# From junk to deleterious: Natural subtelomeric repeat amplifications impact fitness and cellular phenotypes in yeast

**DOI:** 10.64898/2026.06.29.735415

**Authors:** Mathieu Hénault, Virginia Fogg, Lydia R. Heasley

## Abstract

Eukaryotic genomes exhibit astounding levels of complexity. Much of this complexity resides in repetitive DNA thought to evolve neutrally, meaning that its impact on fitness is so small that natural selection cannot act efficiently to favor or purge it. Yet, repetitive DNA greatly facilitates the generation of structural variants (SVs), which fuel evolution with both adaptive and deleterious genetic variation. How SVs involving initially neutral repetitive DNA can bring new evolutionarily meaningful impacts is not well understood, in part because finding and interpreting molecular signatures of these transitions using comparative genomics over long evolutionary timescales is challenging. Here, we document one such transition over a microevolutionary timescale using budding yeast population genomics. We characterize multiple massive amplifications of the Y’ element, a highly polymorphic and dispensable subtelomeric tandem repeat. We uncover extreme structural diversity in Y’ tandem amplifications among near-isogenic strains and show that these amplifications bring a significant fitness cost. We further link Y’ amplifications with transcriptome rewiring, heightened DNA replication stress sensitivity, and DNA damage response activation. Together, our results support a model by which massive subtelomeric tandem amplification pushed a repetitive DNA family outside of effective neutrality to become deleterious.

**Significance Statement:** Extant eukaryotic genomes harbor vast amounts of repetitive DNA elements hypothesized to be non-functional and to have accumulated and persisted through neutral evolutionary processes without substantially affecting organismal fitness. Here, we report that an entire non-functional DNA family evolved out of effective neutrality in the baker’s yeast *S. cerevisiae*. The Y′ elements, a family of non-functional subtelomeric repeats, have repeatedly undergone dramatic tandem amplifications at chromosome ends across independently evolving S. cerevisiae populations. We show that these amplifications confer fitness defects of sufficient magnitude to be considered non-neutral within the population-genetic environment of the species. These amplifications are associated with numerous cellular phenotypes that likely contribute to this fitness cost, including core transcriptome rewiring, heightened sensitivity to DNA replication stress and increased DNA damage.

## Background

The idea that extant genomes harbor large amounts of DNA due to selectively neutral evolutionary processes is long-standing [1]. From a population genetics standpoint, the distribution of many genomic features that contribute to genome size and complexity (e.g. mutation rates, the abundance of introns and selfish genetic elements) is consistent with the degree to which random genetic drift opposes the optimizing power of natural selection [2]. Explaining the existence of so much complexity without invoking an active role for natural selection is a compelling and parsimonious hypothesis. Importantly, it does not imply that these DNA elements are inert within the cell. At least 80% of the human genome is associated with various biochemical signatures (transcription, protein binding, epigenetic modification, etc.) yet under 10% is inferred to be selectively constrained [3,4], which is suggested as a more useful definition of biological function than simply exhibiting biochemical activity [5,6]. This hypothesis also doesn’t imply that a whole non-functional DNA family is bound to remain non-functional. For example, transposable elements (TEs) are primarily selfish genetic parasites [7], but individual transposition events can have deleterious effects for their host [8–10] or be adaptively co-opted [11,12].

Much of the non-functional DNA in eukaryotic genomes shares an important property: its repetitive nature facilitates the generation of new structural variants (SVs) including inversions, translocations, and copy number variations (CNVs) [13–17]. With recent progress in the discovery and genotyping of SVs at population scales [18], SVs are emerging as a ubiquitous source of genetic diversity that explains a large portion of phenotypic variation in populations [19]. It is clear that SVs have the potential to generate phenotypic variation of evolutionary significance, and non-functional repetitive DNA plays a central role in this process. However, understanding the mechanism by which a whole family of non-functional repetitive DNA sequences can escape effective neutrality is challenging, especially when this transition leads to deleterious effects. The inference of weak selection from comparative genomics over long evolutionary timescales is complicated by the challenges of deriving molecular evolution models and distributions of fitness effects tailored to SVs [20,21]. On the other hand, strong selection is expected to rapidly purge deleterious variants, likely driving the causal repeat family to extinction long before it can be observed.

Here, we describe such a transition that recently occurred for a repeat family in the baker’s yeast *Saccharomyces cerevisiae*. Yeast is a uniquely powerful experimental model that integrates population genomics and cell biology, making it possible to track the evolutionary origins of SVs and dissect their impacts on cellular function. We identify recent amplifications of Y’ elements, a dispensable family of subtelomeric tandem repeats and show that these amplifications have a substantial fitness cost within a clade of near-isogenic isolates. We further link Y’ amplifications to multiple cellular phenotypes, including transcriptome remodeling and impaired genome maintenance. Together, our results suggest that within the representative population-genetic environment of yeast, massive tandem amplification led Y’ elements in these strains to transition from being neutral to deleterious.

## Results

### Species-wide CN variation of repeat families

We characterized the species-wide repetitive DNA content variation of *S. cerevisiae* by applying whole-genome sequencing (WGS) to a collection of 336 yeast strains (gift of J. H. McCusker; hereafter the YJM collection), comprising isolates sampled from human clinical (41%), domesticated (35%) and environmental (18%) sources (Table S1). Most isolates in the YJM collection are *S. cerevisiae* (314), while the remainder are *Saccharomyces paradoxus* (7), interspecific *Saccharomyces* hybrids (11), other Saccharomycotina species (3), and a single isolate for which the low WGS data quality prevented species inference. The historical selection of homozygous spore derivatives (HSDs) for WGS has obscured the landscape of SVs in yeast [22]. Our curation of the YJM collection excluded all HSDs based on available annotations, yielding 190 new genomes for strains never sequenced before (112) or previously sequenced by proxy as HSDs (78). To integrate these new genomes with the established population structure of the species, we built a maximum likelihood phylogenetic tree comprising the 314 YJM *S. cerevisiae* strains and 96 reference strains selected at random from the 1011 Genomes Project dataset (1011GP) [23] (Fig. S1A). Our dataset recapitulates the established structure of the species, including ploidy variation and phylogenetic enrichment of clinical isolates (Fig. S1B-C).

We quantified the copy number (CN) per haploid genome for all major repeat families by mapping WGS reads on unique representative copies (Fig. 1A, Fig. S2A, Table S1). One family of subtelomeric repeats, the Y’ element, stands out as the most variable of all repeats (standard deviation, SD: 18.7 copies). The lower end of the distribution comprises 13 environmental strains that have at most one copy of the Y’ element, including three Ecuadorean strains that have none. At the upper end of the distribution, 14 strains show large amplifications (>60 copies). While these outliers belong to multiple genetic clades (Fig. 1B, Fig. S2B), 12 are clinical isolates (Fig. 1C, Fig. S2C), corresponding to a significant enrichment (Fisher’s exact test, *p*-value: 1.1×10^-4^).

**Figure 1.**
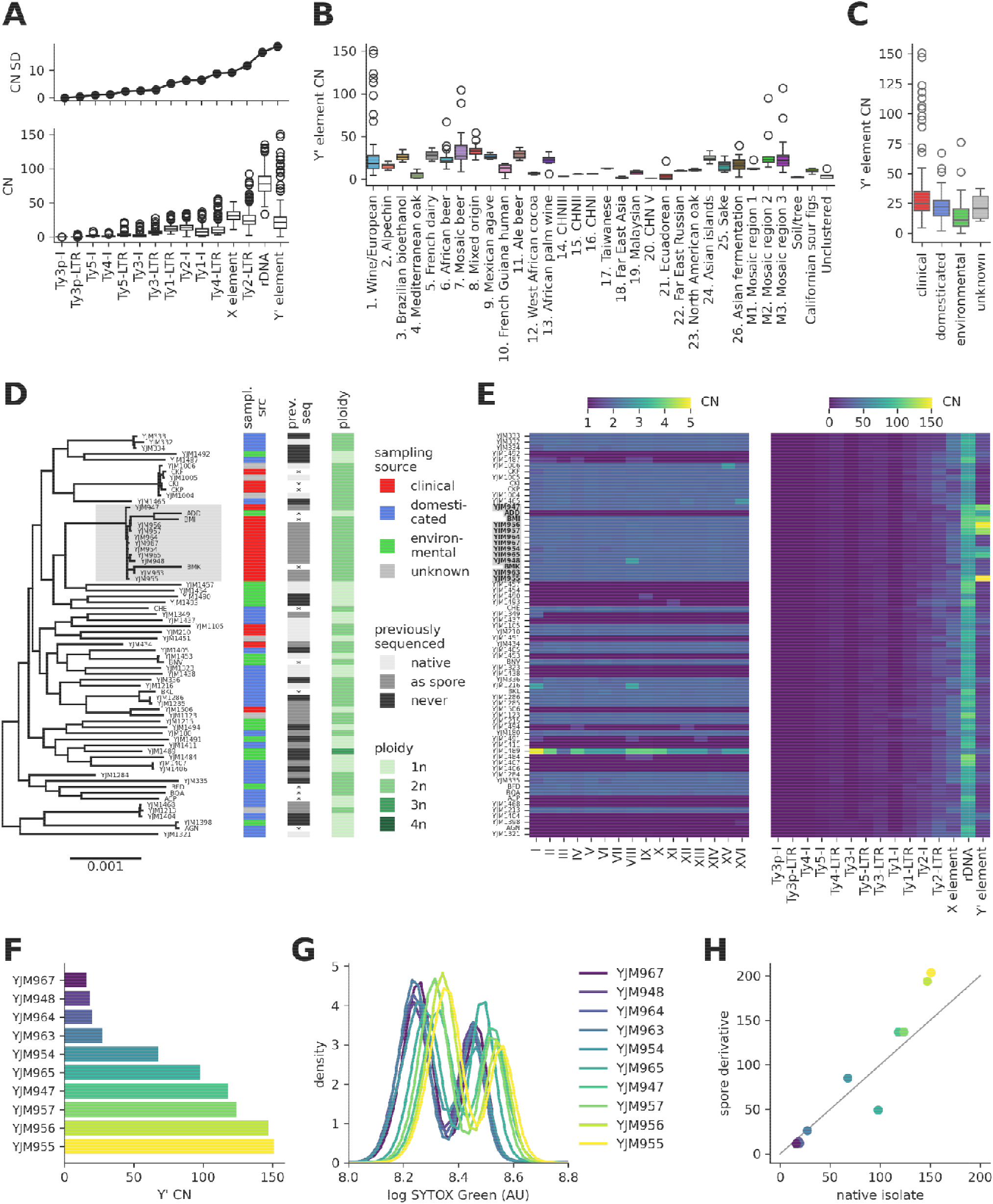
Y’ elements show rapid amplification within a subclade of Wine/European clinical isolates. **A.** Variation in copy number (CN) for major genomic repeat families. Standard deviation (SD) and matching distribution are shown at the top and bottom, respectively. **B.** Distributions Y’ element CN by clade. **C.** Distributions of Y’ element CN by sampling source. **D.** Phylogenetic tree of the Wine/European clade. The wine subclade is highlighted in grey. Annotations from left to right show sampling source, previous sequencing status (asterisks mark the 1011 genomes project [1011GP] strains) and ploidy level measured by flow cytometry. **E.** CN profiles of the 16 chromosomes (aneuploidies, left) and repeat families (right). **F.** Y’ CN distribution for the native strains of the wine subclade. **G.** Distributions of single-cell DNA content of the wine subclade, measured by DNA staining with SYTOX Green and flow cytometry. SYTOX Green fluorescence signal is in arbitrary units (AU). **H.** Y’ CN for the native strains of the wine subclade and their homozygous spore derivatives (HSDs).

Y’ elements are located at subtelomeres, between the X element and the telomere cap, which is composed of TG_1-3_ nucleotide repeats [24]. Typically, Y’ elements are only found at a subset of subtelomeres and can form tandem arrays of up to four copies separated by short interstitial TG_1-3_ repeats [25]. Besides the canonical long (6.7 kb) and short (5.2 kb) variants [26,27], a recent survey detailed much greater Y’ sequence diversity across the species [28]. The Y’ element comprises an ORF, *YRF1*, which encodes a DNA helicase [29], and an autonomously replicating sequence (ARS) near its 3’ end [26]. Tandem Y’ amplifications are adaptive in populations of engineered telomerase-negative mutant cells [29,30]. Otherwise, Y’ elements are completely dispensable for fitness and telomere maintenance [31], an empirically-derived conclusion further corroborated by the existence of multiple natural isolates harboring no Y’ elements. Taken together, these characteristics suggest that Y’ presence/absence polymorphisms are generally neutral and that Y’ elements can be considered non-functional in most natural scenarios.

### Recent Y’ amplification in a subclade of near-isogenic strains

Within the Wine/European clade, one subclade (hereafter referred to as the wine subclade) exhibits a flat population structure, with minimal sequence divergence among strains (excluding three 1011GP HSDs; Fig. 1D, shaded area), and a near-absence of aneuploidy and CNVs for all repeat families, except for the Y’ element (Fig. 1E). The Y’ CN distribution in the wine subclade is hypervariable (SD: 55.4 copies), ranging from 16 to 151 copies per haploid genome (Fig. 1F). Using our previously defined threshold of >60 copies, we categorize six wine subclade strains as harboring clear Y’ amplifications: YJM954, YJM965, YJM947, YJM957, YJM956 and YJM955. Y′ copy number varies by as many as 135 copies among wine subclade strains, corresponding to a ∼6% difference in genome size (∼0.8 Mb). Indeed, the wine subclade strains show a range of single-cell DNA content that correlates with Y’ content variation (Fig. 1G). Using Southern blotting, we observe a corresponding increase in hybridization to a Y’ probe and at least three Y’ size variants that are unevenly distributed among strains (Fig. S3A-C). The striking differences in Y′ content among these otherwise similar genomes suggest substantial and rapid net expansion since their divergence from their last common ancestor.

Y’ amplifications in HSDs of the wine subclade strains have been previously reported [32,33]. De novo assembly of a YJM955 HSD (1011GP ID: ADI) revealed 154 Y’ copies [33], comparable to our estimate for YJM955 (151 copies). While we could not replicate the 1011GP claim that five strains have >150 Y’ copies, our reanalysis of publicly available WGS libraries [34] revealed substantial deviations between native isolates and their HSDs, especially for high-Y’ backgrounds (Fig. 1H). Indeed, we also observe a high deviation in Y’ CN for the only other strain in our collection that harbors a large Y’ amplification (YJM455, 106 copies) and its known HSD (YJM470: 171 copies). We predict that these discrepancies may originate from pervasive heterozygosity in Y’ array length between homologous chromosome ends, leading to increased CN variance upon meiotic segregation. This hypothesis would suggest that Y′ arrays represent an exception to the otherwise low levels of heterozygosity characteristic of the Wine/European clade (Fig. S1A).

### Extreme structural diversity in Y’ amplifications

To gain insights into the structural organization of Y’ amplifications, we performed Oxford Nanopore (ONT) long read sequencing for 48 YJM strains, including the whole wine subclade and six strains with amplifications outside of the Wine/European clade. We annotated Y’ sequences from our unassembled ONT reads and the genome assemblies of the *Saccharomyces cerevisiae* Reference Assembly Panel (ScRAP) [33], covering the complete phylogenetic diversity of the species (Fig. S3D). Using the 22381 annotated Y’ sequences originating from both sources (see Methods), we employed agglomerative clustering to define 17 main Y’ consensus types (Fig. S4), which are distinguishable primarily by internal and terminal deletions (Fig. S3E). In agreement with a recent report of species-wide Y’ sequence diversity [28], a single type is exclusive to the wine subclade and responsible for the largest amplifications (type 10; Fig. S3F) [25]. However, other amplifications across the phylogeny contain different Y’ types, suggesting that most Y’ sequence variants can fuel large-scale subtelomeric expansions.

In addition to phasing individual Y’ copies, sufficiently long reads can also resolve tandem organization at the haplotype level. Because this requires a high coverage of ultra-long reads, we focused specifically on the wine subclade. Haplotype-aware de novo assembly algorithms consistently failed to phase contigs all the way to telomere caps. To circumvent this, we anchored raw reads to unique sequences at each chromosome end, visualized their Y’ type annotations and manually extracted the consensus structure for all major haplotypes (Fig. 2). The conservation of unique centromere-proximal gene and element sequences across all reads in the alignments to each chromosome end validates the specificity of this approach. Across the wine subclade, Y’ arrays display remarkable heterozygosity, with two distinct haplotypes found at most homologous chromosome ends (Fig. 2). Arrays are very heterogeneous, containing at least two Y’ types often present in complex patterns. While some telomere caps are missing from our analysis (likely an artefact of limited read lengths), we find that Y’ amplifications are associated with longer telomere caps (Fig. S5A-B), consistent with previous studies [35,36]. Interstitial TG_1-3_ repeats within Y’ amplifications show a bimodal length distribution (Fig. S5C), a likely consequence of their association with specific Y’ types (Fig. S5D). These data suggest that Y’ amplifications are very dynamic and quickly generate new structural diversity.

**Figure 2.**
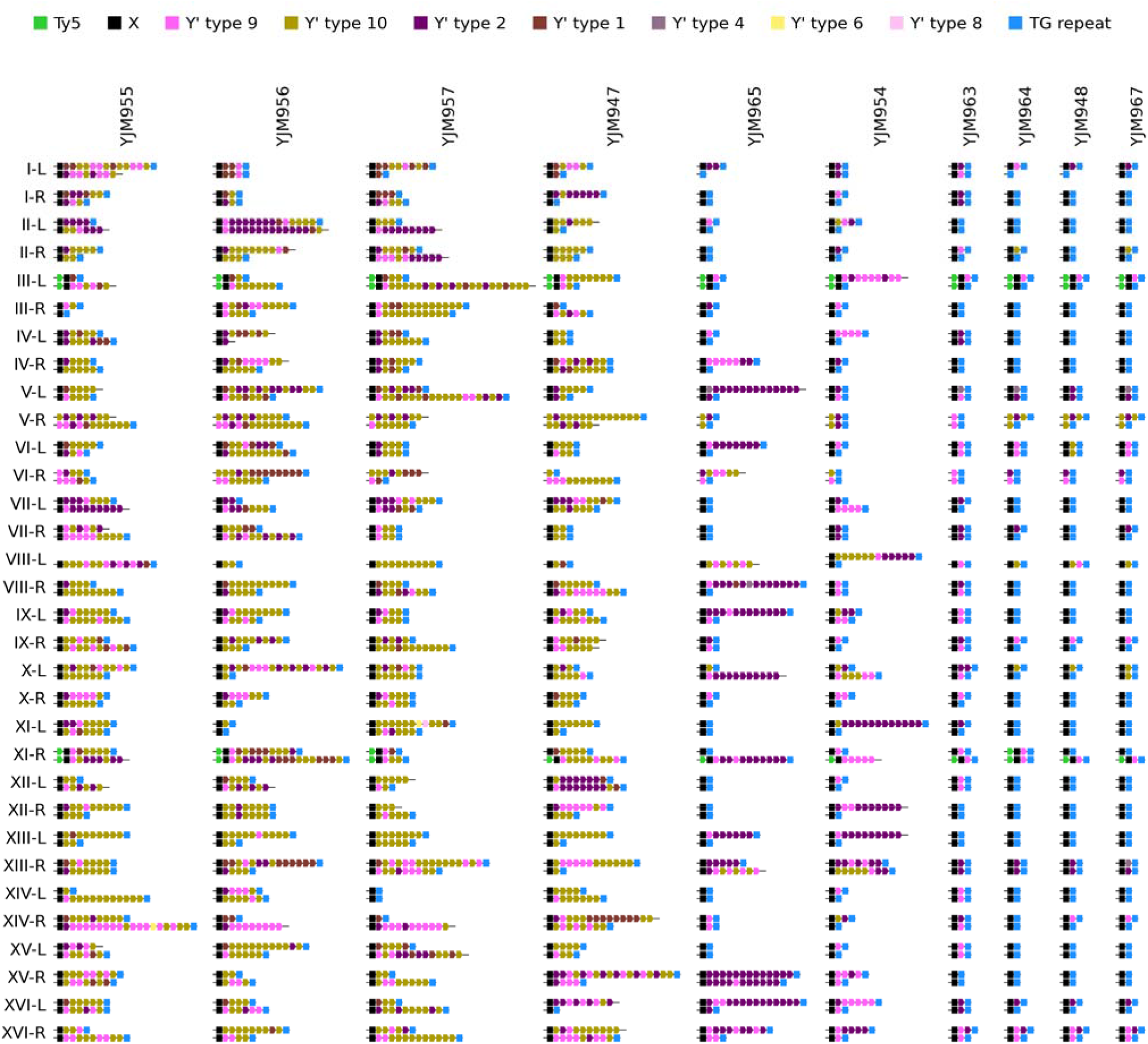
Y’ amplifications display pervasive heterozygosity and complex heterogeneity in the wine subclade. For each chromosome end, symbols show the elements found in the two major subtelomeric haplotypes, from Ty5/X at the left to the telomere cap at the right. Haplotypes without a cap are deemed incompletely resolved and marked by an extended thin line.

### Fitness cost of Y’ amplifications

Carrying Y’ amplifications could be costly for the cell, for example by imposing additional gene expression, DNA replication, and/or DNA damage repair burdens. We tested this hypothesis by measuring growth kinetics of the wine subclade strains in permissive conditions, in both complex (YPD) or defined (SC) media. We find a significant negative correlation between maximum growth rate and Y’ content (YPD, R^2^=0.779, p=7.2×10^-4^; SC, R^2^=0.899, p=3.0×10^-5^), corresponding to a selection coefficient of 6.9 to 9.5 % (Fig. 3A). We considered the possibility that other genomic variants may independently contribute to growth rate or co-vary with Y′ content across strains. To test this, we performed a genome-wide association study (GWAS) to identify core genome variants associated with maximum growth rate in YPD. No other variant exceeds the threshold for statistical significance defined by permutation analysis (Fig. 3B). Many of the top GWAS scores are variants at the very ends of the chromosomes, and thus in strong linkage disequilibrium with the neighboring Y’ array. Only rDNA CN is significantly positively correlated with Y’ content in the wine subclade (Fig. 3C). However, the correlation of rDNA CN with growth rate is much weaker than for Y’ CN(Fig. 3D) and previous data indicate that a difference of ∼50 rDNA copies in laboratory strains has a fitness cost of ∼0.5% [37,38], an order of magnitude lower than our estimate. Together, these data indicate that Y’ content is the best genetic predictor of maximum growth rate variance in permissive conditions. These data suggest that while Y’ amplifications are likely not the only cause of the quantitative fitness reduction within the wine subclade, they are the greatest contributor.

**Figure 3.**
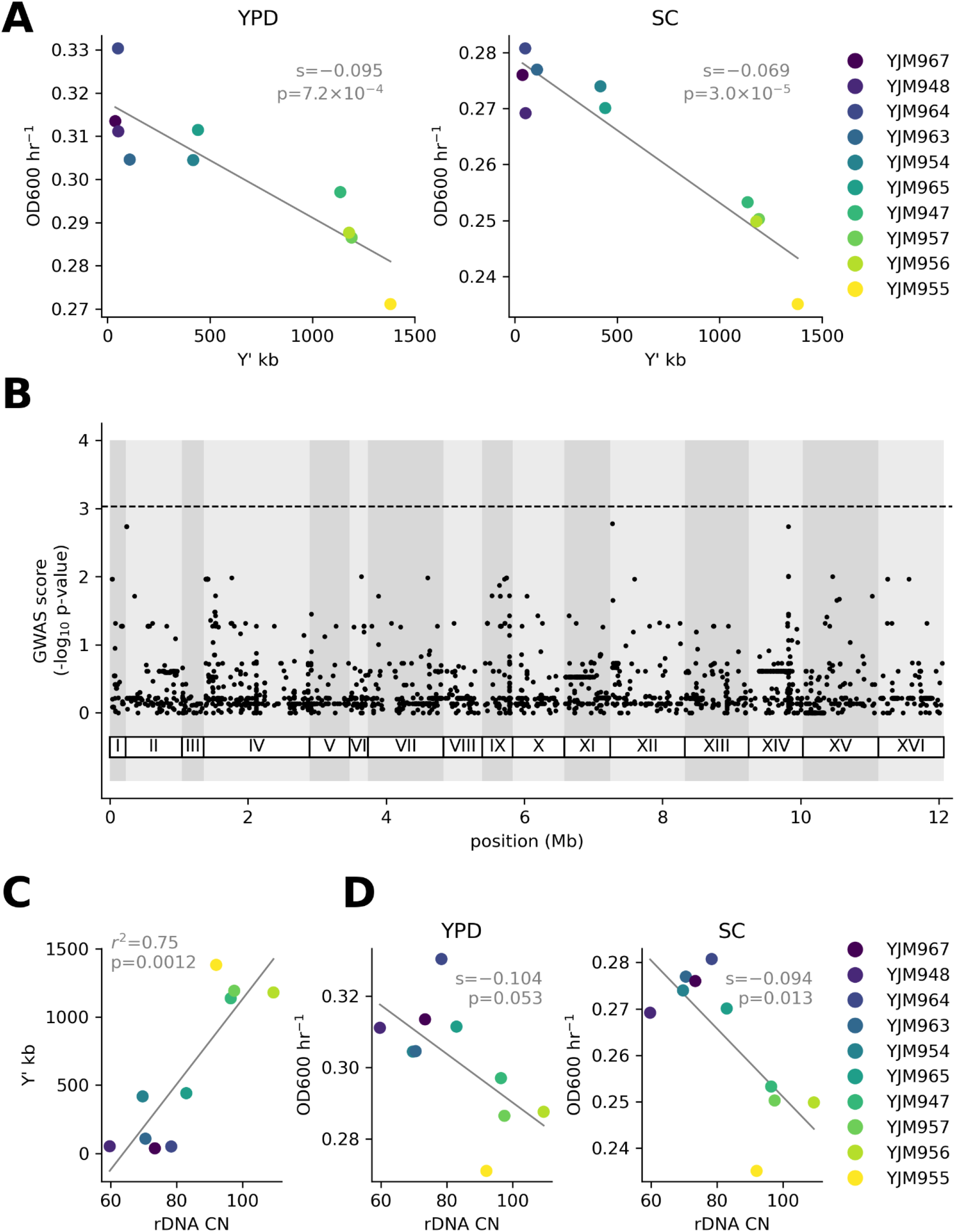
Y’ content is linked to a gradual fitness cost. **A.** Maximum growth rate extracted from growth curves in liquid rich medium, either complex (YPD) or defined (SC), measured as optical density at 600 nm (OD600). P-values for Pearson’s correlation against Y’ content, as quantified by long-read annotations, are shown. The Malthusian selection coefficient (*s*) was calculated as the ratio of fitted maximum doubling rates at the extremes of the Y’ content range (38 vs 1382 kb). **B.** Genome-wide association study (GWAS) for maximum growth rate in YPD. The dashed line corresponds to the significance threshold determined by permutation analysis. **C.** Correlation of Y’ CN against rDNA CN. P-value and r^2^ are shown for Pearson’s correlation. **D.** Correlation of maximum growth rate as in A, but against rDNA CN.

### Y’-associated transcriptome remodeling

We next investigated the molecular basis of the fitness defect associated with Y’ amplifications. We profiled the transcriptomes of four strains along the Y’ CN spectrum. We find 373 significant differentially expressed (DE) genes whose abundance changes a function of increasing Y’ content, both upregulated (158) and downregulated (215; Fig. 4A). The Y’ element stands out as the largest positive change, with a ∼10 fold increase between low and high-Y’ backgrounds (Fig. 4A-B, Fig. 1F). This increase is consistent with individual Y’ copies being expressed across backgrounds. As a result, the Y’ transcript ranks among the most abundant in high-Y’ strains, accounting for ∼1% of the total mRNA in the cell, implicating Y’ transcriptional burdens as an appreciable component of the observed fitness cost.

**Figure 4.**
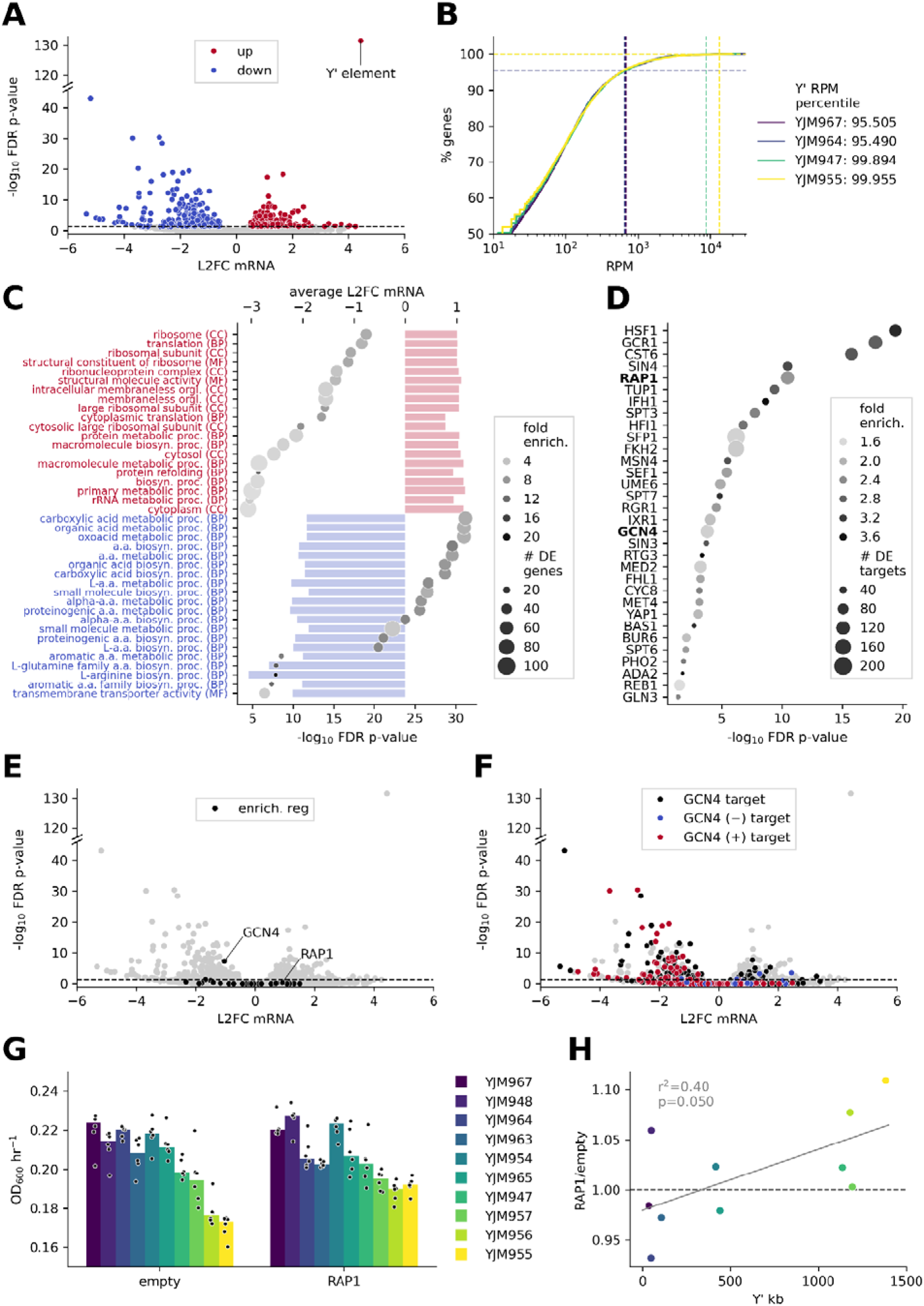
Transcriptome-wide signatures are associated with Y’ amplifications. **A.** Differentially expressed (DE) levels as a function of Y’ content. Upregulated and downregulated genes are shown in red and blue, respectively. The dashed line corresponds to the DE significance threshold of 0.05. **B.** Cumulative distributions of the top 50% transcript levels in reads per million (RPM). For each gene, RPM values are averaged across the three replicate libraries, and the resulting average RPM values are sorted by ascending order along the X axis, while their corresponding percentile is plotted on the Y axis. Dashed lines show the RPM and percentile of the Y’ element in each transcriptome. **C.** Gene ontology (GO) terms enriched among the significantly DE genes as a function of Y’ CN. Dot position on the X axis corresponds to the significance level of the enrichment (P-value). Dot size and greyscale value indicate the number of DE genes annotated with the given GO term and associated fold enrichment, respectively. Bars indicate the average log_2_ fold change (L2FC) for the DE genes annotated with the corresponding GO term. The 20 most significantly enriched (lowest P-values) up and downregulated terms are shown. BP: biological process, CC: cellular compartment, MF: molecular function. **D.** Regulators whose targets are significantly enriched among DE genes. Dot size and greyscale value indicate the number of DE genes annotated as targets of the given regulator and associated fold enrichment, respectively. FDR-adjusted p-values for Fisher’s exact tests are shown. **E.** Same as A, but highlighting the regulators whose targets are significantly enriched among DE genes. **F.** Same as A, but highlighting the regulatory targets of *GCN4*, and whether the sign of the regulation by *GCN4* is positive (red), negative (blue) or unannotated (black). **G.** Maximum growth rate of the wine subclade transformed with an empty vector plasmid (left) or a vector expressing *RAP1* (right). **H.** Ratio of maximum growth rate for cells carrying the *RAP1* overexpression plasmid to the empty plasmid control. Pearson’s correlation is shown.

We next probed how Y’ abundance modulates core genome expression. Ribosome and carboxylic acid metabolic processes are the most significantly upregulated and downregulated gene ontology (GO) terms, respectively (Fig. 4C). Integrating the yeast regulatory network with these data, we identified 32 transcriptional regulators with targets that are significantly overrepresented among DE genes (Fig. 4D). The only significantly enriched regulator that is itself also significantly DE is the general amino acid control master effector *GCN4*, with a 2.05 fold decrease in transcript level (Fig. 4E). As predicted, all 43 differentially expressed (DE) targets positively regulated by Gcn4 are downregulated, while 3 of the 5 DE targets negatively regulated by Gcn4 are upregulated (Fig. 4F). This result indicates that the regulation of multiple core metabolic processes, including amino acid biogenesis, is altered in the presence of Y’ amplifications.

Our regulatory network analysis highlighted *RAP1* as the fifth most significantly enriched regulator, with 119 DE targets (Fisher’s exact test, FDR-adjusted *p*-value: 3.5×10^-11^). Unlike *GCN4*, the transcript level of *RAP1* itself does not change significantly as a function of Y’ content (Fig. 4E). However, interstitial TG_1-3_ repeats within Y’ arrays are strong Rap1 binding sites [39], prompting the hypothesis that the increased number of subtelomeric binding sites in strains with Y′ amplifications could deplete the Rap1 pool available for proper regulation of its core genome targets, leading to a reduction in growth rate. If this model is correct, increasing *RAP1* expression should restore Rap1 availability at its core genome targets and thereby mitigate the growth defect associated with Y′ amplifications. We tested this by overexpressing *RAP1* from a high-copy plasmid [40]. Cells carrying an empty vector control exhibit the same correlation between growth rate and Y’ content as WT cells (Fig. 4G). When overexpressing *RAP1*, high-Y’ strains specifically show an increase in growth rate (Fig. 4H, Pearson’s correlation *p*-value: 0.050), indicating that *RAP1* overexpression partially rescues the fitness defect linked to Y’ amplifications. This result supports a model in which Y’ amplifications can preferentially sink sequence specific regulatory factors like Rap1p away from their regulatory targets.

### Y’-associated vulnerability in genome maintenance

We next asked whether Y’ amplifications are a burden on the replication of the genome. Net gain aneuploidy has been linked to increased DNA replication stress sensitivity in mammalian cells [41,42], and we hypothesized that Y’ amplifications could cause a similar effect. The wine subclade comprises near-isogenic strains spanning a gradient of Y′ content, with one strain carrying an additional chromosome VIII trisomy (Fig. 5A) providing a natural system to assess the relative contributions of Y′ amplification and aneuploidy to replication stress sensitivity. We quantified the growth kinetics of cells exposed to hydroxyurea (HU), a compound that challenges DNA replication by depleting intracellular dNTP pools [43]. HU addition magnifies the Y’-associated defect measured in permissive conditions (Fig. 5B). From dose-response curves in HU (Fig. 5C), we extracted the half-maximal inhibitory concentration (IC50) as a quantitative measure of inhibition by HU (Fig. 5D). There is a strong correlation between extra DNA content (either Y’ or VIII) and HU IC50 (Fig. 5E). This result indicates that extra genomic DNA, be it in the form of Y’ amplifications or aneuploidies, increases DNA replication stress sensitivity.

**Figure 5.**
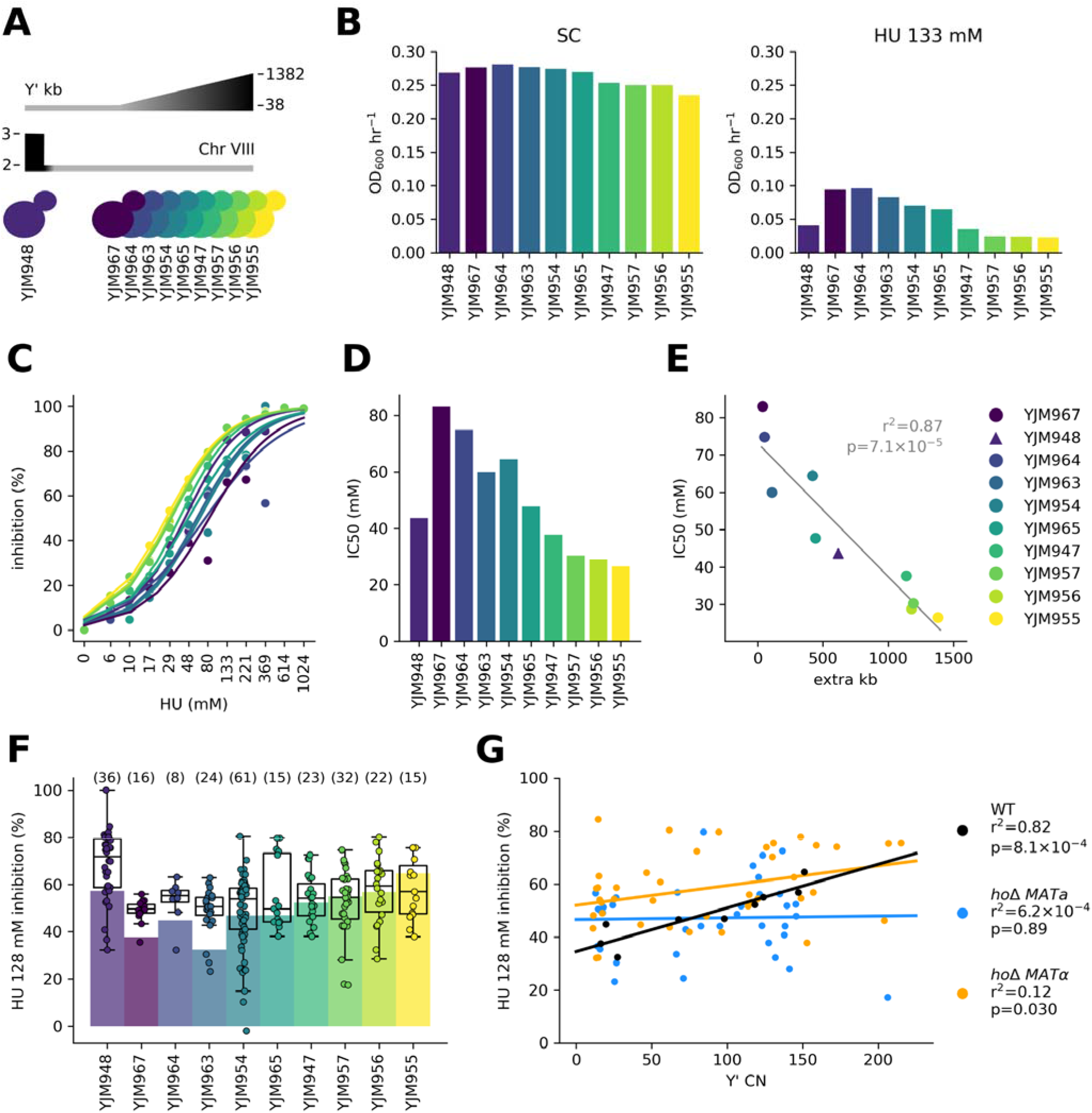
Y’ amplifications correlate with DNA replication stress sensitivity. **A.** The wine subclade as a model system to assess the impact of increased DNA contents, both in the form of net gain aneuploidy (VIII trisomy in YJM948) and Y’ amplification, in otherwise nearly isogenic background . **B.** Maximum growth rate of the wine subclade in SC and 128 mM HU. **C.** Dose-response curves of the wine subclade in HU. **D.** Half-maximal inhibitory concentration (IC50) of the wine subclade. **E.** Correlation between HU IC50 and extra DNA content, either in the form of Y’ amplification or VIII trisomy. **F.** Growth inhibition of haploid spores derived from each background of the wine subclade, measured as the ratio between growth rate in 128 mM HU and SC. Bars indicate the inhibition level of the diploid backgrounds. **G.** Pearson’s correlation between HU inhibition and Y’ CN for WT diploids and MAT**a** or MATα haploid spores.

To further probe the link between DNA content and HU sensitivity, we generated a collection of HSDs for each strain of the wine subclade, reasoning that meiotic segregation would generate HSDs harboring a broader and more finely resolved range of net Y′ content, allowing us to more precisely define the relationship between Y′ CN and HU sensitivity. Sequencing of 84 HSDs confirmed the expanded range of Y′ CN within each background and across the whole subclade. Sequencing also revealed that Y’ CN variance is greater in high-Y’ backgrounds harboring the most pronounced levels of heterozygosity between homologous arrays (Fig. S6A). Unexpectedly, however, we find that the correlation between Y’ content and HU sensitivity observed in the diploid parental strains completely vanishes in HSDs, both within individual parental backgrounds and across the wine subclade (Fig. 5F-G). Quantitative trait loci (QTL) analysis revealed several SNVs associated with HU sensitivity (Fig. S7A), although with modest effect sizes (Fig. S7B). Inherited aneuploidy of VIII has a marginally significant but larger effect (n=10, Student’s *t*-test, *p*-value: 0.08, Fig. S7B). We reasoned that two haploid-specific features could explain the loss of Y’ CN-associated HU sensitivity in HSDs: 1) they harbor half the number of chromosomes relative to the diploid parental strains, or 2) they express a single mating type (*MAT*). To distinguish between these possibilities, we made hemizygous deletions of the *MAT* locus in the parental diploid strains such that only one mating type was expressed. We observe no change in HU sensitivity (Fig. S8). Thus, our data show that DNA replication stress sensitivity associated with Y’ content and aneuploidy are respectively dependent and independent of base ploidy level. Whether the diploid-specific Y’-associated HU sensitivity is caused by total cellular DNA content or the possible involvement of homolog-homolog interactions [44] remains to be tested.

Y’ elements are euchromatic [45], transcribed [46], and late replicating [47]. Transcription-replication collisions can lead to DNA breaks at stalled replication forks [48], and stalled forks are exacerbated by HU exposure [49]. To validate that Y’s are transcribed upon HU exposure, we augmented our transcriptome dataset with strain samples grown at their respective HU IC25, IC50 and IC75 (Fig. S9A-B). Using published gene expression measurements from the yeast GFP collection in 200 mM HU [50], we validated that HU exposure in our transcriptomes recapitulates known transcriptional responses, including overexpression of the ribonucleotide reductase inhibitor *HUG1*, which occurs in response to DNA damage [51–53]. Y’ transcript levels exhibit a modest increase with HU exposure level (Fig. S9C-D). We next determined whether Y’ amplifications are associated with increased DNA damage by quantifying the frequency of *RAD52* foci [54]. Using the same four backgrounds as the transcriptomics dataset, we tagged one allele of *RAD52* with a fluorescent reporter and imaged cells grown in SC or at their respective HU IC50. Even in the absence of HU, cells from high-Y’ backgrounds more frequently exhibit *RAD52* foci and, when they do, tend to have multiple foci per cell (Fig. 6A-C). Both Y’ content and HU exposure have statistically significant effects on the presence and abundance of foci per cell (Fig. 6D). Our microscopy data also suggested that high-Y’ strains have reduced viability in HU, which we validated by spot assays on solid medium (Fig. S10). These results indicate that Y’ amplifications are associated with DNA damage, and this effect is stronger upon DNA replication stress. Whether Y’ arrays themselves are local DNA damage hotspots, perhaps as a consequence of transcription-replication conflicts, requires further investigation.

**Figure 6.**
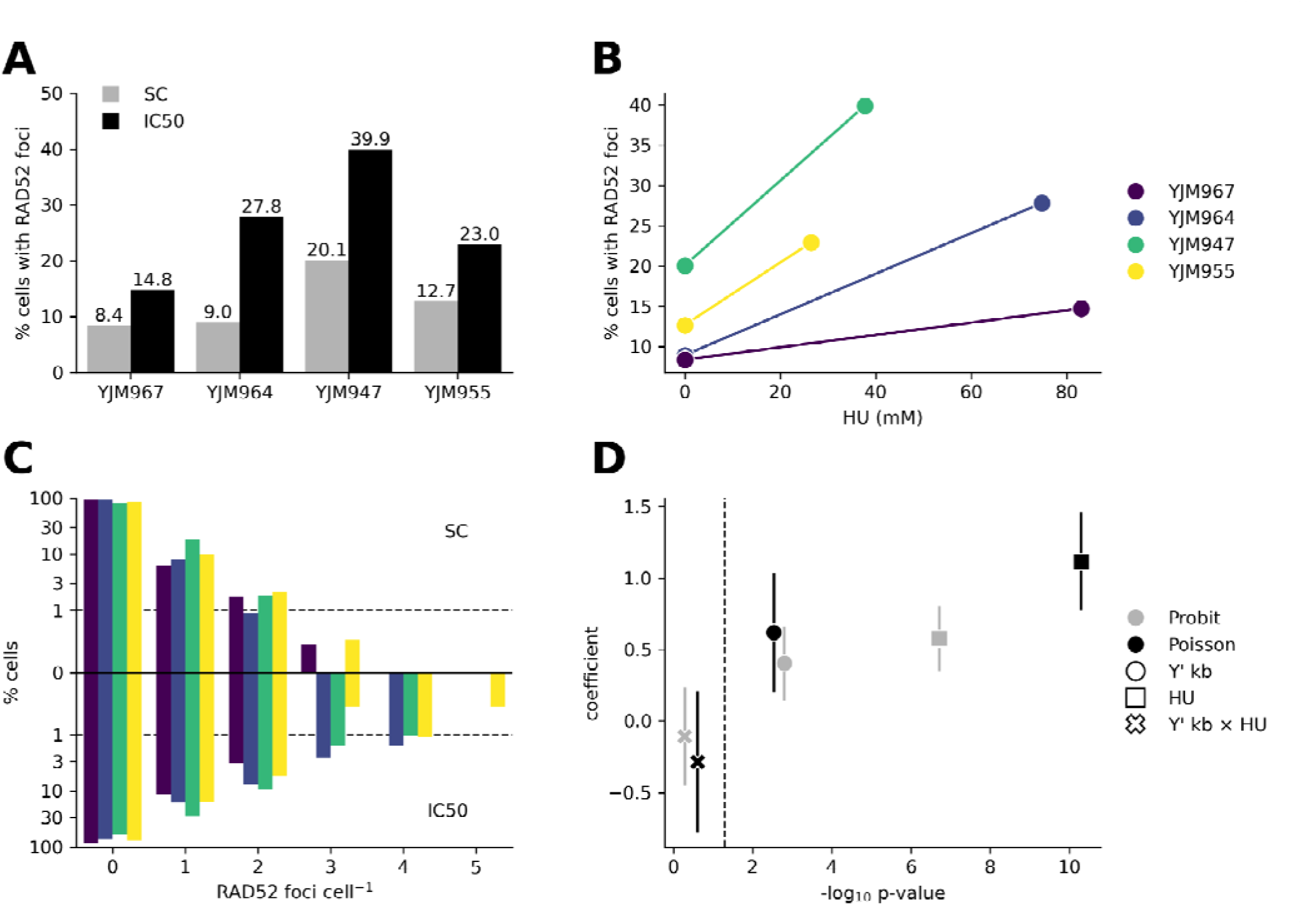
Amplified Y’ elements are linked to increased DNA damage response. **A.** Frequency of cells with *RAD52* foci, measured by confocal microscopy in SC and at HU IC50. **B.** Frequency of cells with *RAD52* foci as a function of HU dose. **C.** Distributions of *RAD52* foci counts per cell in SC (upper) and HU IC50 (lower). Symlog transformed frequencies are plotted, and dashed lines correspond to the 1% threshold for log transformation. **D.** Coefficient and *p*-value for terms of generalized linear models fitted on the presence (Probit) and count (Poisson) of *RAD52* foci per cell. The dashed line corresponds to the significance threshold of 0.05.

## Discussion

Understanding the mechanisms and dynamics by which evolutionary forces shape genome sequences is a central goal of biological research. Here, we uncover a family of subtelomeric repetitive elements that transitioned from being selectively neutral to being deleterious through massive tandem amplification. This transition appears to differ from how most non-functional repetitive DNA is thought to become subject to natural selection: rather than a few repeats acquiring selective effects, such as individual TE copies disrupting protein-coding sequences, our data suggest a mass effect in which increasing cumulative Y′ content progressively imposes a stronger fitness cost.

The natural Y’ amplifications we have characterized here are reminiscent of laboratory generated mutants that display an alternative lengthening of telomeres (ALT) phenotype, specifically type I survivors that arise in populations of various telomerase mutant cells [30]. In these conditions, Y’ amplifications are strongly beneficial since they rescue the erosion of chromosome ends that happens after at least 30 mitotic divisions [36,55,56]. We emphasize that multiple features of the subtelomeric architectures harbored by the wine subclade strains are inconsistent with a type I survivor phenotype. First, they exhibit telomere caps of several hundreds of base pairs in length, incompatible with cap erosion. Second, the high level of Y’ sequence heterogeneity within individual arrays is in stark contrast with type I survivors, which exhibit mostly homogeneous arrays [36]. Third, no gene known to be involved in telomere maintenance harbors variants with a high predicted functional impact in the wine subclade. We cannot exclude that by an unknown mechanism, these strains experienced transient telomerase dysfunction, which was recently shown to be sufficient to initiate Y’ expansions [57]. However, we argue that this is an unlikely scenario for wild type diploid natural isolates, since all documented forms of ALT involve a complete loss of function of at least one gene (e.g. *EST1*, *TLC1* or *CDC13*) in a haploid context.

The evolutionary path by which the Y’ amplifications harbored by these extant wild isolates arose is unclear. We uncover a strong association with human clinical isolates [58], and one or multiple components of this environment (temperature, pH, carbon source) could impose a biased mutation pressure towards the expansion of Y’ arrays. For example, exposure to high temperatures during opportunistic infection in humans could drive subtelomeric instability by triggering the inherent thermolability of telomeres [59]. A non-mutually exclusive hypothesis is that Y’ amplifications are adaptive in opportunistic infections. Passage through a host likely alters selection pressures while also magnifying the power of random genetic drift through reduced effective population size (*N_e_*) [60]. A population bottleneck implies that the selection coefficient (*s*) linked to Y’ amplifications was necessarily large to escape effective neutrality (i.e. |*N_e_s*| > 1) [61]. Although unlikely for the reasons discussed above, a large *s* is expected for type I survivors arising from a persistent telomerase-deficient background. We also cannot exclude that Y’ amplifications were driven to near-fixation by drift alone.

Regardless of the conditions that permitted their emergence, Y’ amplifications show a substantial fitness cost in permissive conditions, with an estimated *s* of -0.05 to -0.10. Most estimates of *N_e_* for *S. cerevisiae* are in the 10^6^-10^7^ range [62,63], meaning that both *N_e_* and *s* would need to be overestimated by roughly two orders of magnitude for the Y’ amplifications carried by these strains to be compatible with effective neutrality at the species level (i.e. |*N_e_s*| < 1).

In addition to the population genetic environment that permitted Y’ amplifications to arise, the mutational mechanisms underpinning their development and maintenance are also currently unknown. Y’ arrays within the wine subclade display an enormous degree of heterogeneity, suggesting that these loci are inherently structurally dynamic. The tandemly repeated and homotypic architecture of yeast subtelomeres undoubtedly contributes to this plasticity. Stochastic nonallelic homologous recombination (NAHR) occurring within and between Y’ arrays can generate new CNVs [64], but how variation in the copy number and distribution of Y’ elements across the chromosome ends impact the rates and outcomes of subtelomeric NAHR have not been investigated. We predict that rates of subtelomeric NAHR should increase as a function of total Y’ element copy number, with the magnitude of NAHR-mediated amplifications or deletions also depending on the heterogeneity of Y’ array lengths across chromosome ends. The wine subclade strain group provides a unique experimental system with which to investigate these open questions in the future and to determine how mutational processes drive the evolution of non-functional repetitive sequence content in genomes.

In conclusion, Y’ elements show the hallmarks of repetitive non-functional DNA, being dispensable for organismal fitness and having no demonstrated function from an evolutionary standpoint. We show that large tandem Y’ amplifications arose as extant natural polymorphisms that are detrimental to organismal fitness. We suggest that in this specific context and within the representative long-term population-genetic parameters of the species, they no longer fit the view of near-neutral genetic hitchhikers.

## Methods

### Strains and media

Strains used in this study are listed in Table S1-3. Culture media were prepared as follows. YPD: 10 g L^-1^ yeast extract (US Biological cat. Y2010), 20 g L^-1^ peptone Y (US Biological cat. P3306), 20 g L^-1^ glucose (Sigma Aldrich cat. G7021). SC: 1.72 g L^-1^ yeast nitrogen base (US Biological cat. Y2030), 2.06 g L^-1^ complete drop-out (US Biological cat. D9516), 20 g L^-1^ glucose, 5 g L^-1^ ammonium sulfate (Fisher Scientific cat. BP212R-1). SC-MSG: 1.72 g L^-1^ yeast nitrogen base, 2.06 g L^-1^ complete drop-out, 20 g L^-1^ glucose, 1 g L^-1^ L-Glutamic acid monosodium salt (MP Biomedicals cat. 101800). SPO: 10 g L^-1^ potassium acetate (Thermo Scientific cat. A16321.36), 1 g L^-1^ yeast extract, 0.5 g L^-1^ glucose, 0.35 g L^-1^ complete drop-out. 2YT: 10 g L^-1^ yeast extract, 16 g L^-1^ peptone Y, 2 g L^-1^ glucose, 5 g L^-1^ sodium chloride (Fisher Scientific cat. BP358-212). Solid media plates were prepared by adding 20 g L^-1^ agar (US Biological cat. A0930). G418 sulfate (GoldBio cat. G-418-25) was supplemented at 200 mg L^-1^. Hygromycin B (GoldBio cat. H-270-10) was supplemented at 250 mg L^-1^. Nourseothricin sulfate (GoldBio cat. N-500-1) was supplemented at 100 mg L^-1^. Ampicillin sodium (GoldBio cat. A-301-5) was supplemented at 100 mg L^-1^.

### Molecular cloning

Primers are listed in Table S4. PCRs for the generation of gene deletion cassettes, plasmid fragments or hybridization probes were done using Q5 high-fidelity polymerase (NEB cat. M0491) or KAPA Hifi Hotstart Readymix (Roche cat. KK2602). Site directed mutagenesis was performed with the KAPA Hifi Hotstart Readymix. Molecular assembly was performed using the NEBuilder HiFi DNA Assembly Master Mix (NEB cat. 2621). Transformations were done in *Escherichia coli* strain DH5α using a standard CaCl_2_ protocol. Plasmid minipreps were done on cells grown in 2YT+Amp using the GeneJET Plasmid Miniprep Kit (Thermo Scientific cat. K0503). Transformations in yeast were done using a standard LiOAc-PEG protocol. Confirmation PCRs were run with GoTaq G2 Green master mix (Promega cat. M7823).

### Single-cell DNA content measurement

Cells from saturated precultures were resuspended in sterile water, fixed in 70% ethanol and treated overnight with RNAse A (Fisher Scientific cat. BP2539) at 37°C in sodium citrate buffer (50 mM, pH 7.0). Cells were stained in 0.6 μM SYTOX Green (Invitrogen cat. S7020) and analyzed on a ZE5 YETI flow cytometer (Bio-Rad). Cytometry data were analyzed using FlowKit v1.3.0 [65] in a custom Python v3.12 script [66].

### Reference genomes

Unless otherwise stated, the reference genome used is S288C vR64-4-1_20230830 (https://www.yeastgenome.org). A masked reference genome with unique repeat family sequences appended as artificial contigs was generated by replacing all repeats and regions of uneven short-read coverage at chromosome ends with Ns. Family consensus sequences for X elements, Y’ elements and rDNA units were generated as follows. Annotated sequences were extracted from genome assemblies of strains S288C, DBVPG6044, SK1, YPS128 and UWOPS03-461.4 [67] using a custom Python script. Sequences were aligned using Muscle v5.1[68] and alignments were trimmed using trimAl v1.4 [69] in gappyout mode. Consensus sequences were computed using the cons tool from the EMBOSS v6.6.0.0 suite [70]. TG repeats of the S288C reference genome were extracted and concatenated as a single artificial contig. Ty element reference sequences were taken from Carr et al. [71].

### Whole-genome sequencing and variant calling

Original stocks were streaked for single colonies on YPD agar plates and one single colony was picked to inoculate a 5 mL YPD culture grown overnight at 30°C. Genomic DNA was extracted using the YeaStar Genomic DNA Kit (Zymo Research cat. D2002). Libraries were prepared using the plexWell 384 kit (seqWell cat. PW384) and sequenced on one lane of a 10B or 25B NovaSeq X Plus flow cell (Illumina) in paired-end 150 layout (Novogene). Reads were trimmed using fastp v1.1.0 [72] with default parameters and aligned to the masked reference genome version using bwa-mem2 v2.2.1 [73]. Secondary alignments were excluded using the view tool from the samtools v1.23 suite [74]. Read duplicates were marked and excluded using the MarkDuplicates tool from the picard v3.1.1 suite (http://broadinstitute.github.io/picard), and read group tags were added using picard AddOrReplaceReadGroups. Variants were called using the mpileup tool from the bcftools v1.19 suite [75] and bcftools call with options -m -v. Variant tags were added using the bcftools plugin fill-tags and erroneous heterozygous calls (variant allele frequency <0.15 or >0.85) were corrected using the bcftools plugin setGT. Variants with quality scores below 20 were excluded.

### Phylogenetic and population structure analysis

We build a maximum likelihood tree for 326 YJM strains and 96 reference strains from the 1011GP [23]. We selected 500 ORFs at random, computed their consensus sequence with bcftools consensus and concatenated them using a custom Python script. A maximum likelihood phylogenetic tree was computed using raxml-ng v1.2.2 [76] with model GTGTR4 and 200 bootstrap trees, specifying Taiwanese isolates AMH, CEG and CEI as outgroups [23]. Clade was assigned to each isolate of the YJM collection based on placement in the tree in relation to the 1011GP strains and the clade previously assigned to each of the latter [23]. A BED (PLINK binary biallelic genotype table) file was generated from the calls in VCF format using PLINK v1.9.0 [77], filtering for linkage disequilibrium in 50 kb windows with steps of 10 variants and an r^2^ threshold of 0.1. Population structure was computed using ADMIXTURE v1.3.0 [78] with 5-fold cross-validation for K values of 3 to 41. The optimum K value determined by cross-validation is 29.

### Long read sequencing

High molecular weight (HMW) genomic DNA was prepared from 5 mL overnight YPD cultures. Cells were digested with zymolyase 20T overnight at 37°C and lysed in SDS, followed either by cell lysate precipitation in KOAc and DNA precipitation in isopropanol, or double phenol-chloroform-isoamyl alcohol extraction and DNA precipitation in ethanol. RNA was digested with RNAse A at 37°C. Multiplex native libraries were prepared using the SQK-NBD114-24 kit (Oxford Nanopore Technologies). Libraries were either sequenced with FLO-PRO114M flow cells on a PromethION 2 Solo sequencer (ONT), or with FLO-MIN114 flow cells on a MinION Mk1B sequencer. Basecalling was run at super high accuracy using dorado v0.7.2 (ONT).

### Southern blot

A 418 nt probe corresponding to the conserved central region of Y’ elements (Fig. S3A) was amplified from YJM955 genomic DNA with primers MHO_0035 and MHO_0036 and labelled with the ULYSIS Alexa Fluor 488 Nucleic Acid Labeling Kit (Invitrogen cat. U21650). For each strain of the wine subclade, 1 μg of HMW genomic DNA was digested with XhoI (NEB cat. R0146) overnight at 37°C and separated by 0.8% agarose gel electrophoresis. The gel was stained with ethidium bromide and imaged, then nicked in 0.25 N HCl for 20 min and denatured in transfer buffer (0.4 N NaOH, 1M NaCl) twice for 20 min. Alkaline transfer to a BrightStar-Plus Positively Charged Nylon Membrane (Invitrogen cat. AM10104) was done in transfer buffer using a Model 785 Vacuum Blotter (Bio-Rad) at 5 in. Hg for 3 hrs. The membrane was blocked at 42°C for 30 min with ULTRAhyb Ultrasensitive Hybridization Buffer (Invitrogen cat. AM8670) preheated to 68°C. Hybridization was done in fresh hybridization buffer at 42°C overnight. Imaging was done with a Sapphire Biomolecular Imager (Azure Biosystems).

### Y’ types annotation and clustering

Y’ element sequences were annotated and extracted as follows. Assemblies from the ScRAP panel [33] were aligned against the masked S288C reference using minimap2 v2.28 [79] with preset asm10. From the basecalled ONT libraries, the longest reads corresponding to 5X genome coverage (60 Mb, minimum length: 1 bp, maximum length: 656189 bp, average length: 27128.3 bp) were subset using a custom Python script and aligned to the masked S288C reference using minimap2 with preset lr:hq. Alignments to the Y’ consensus in PAF format were converted to BED format using a custom Python script and sequences in FASTA format were extracted using bcftools v2.31.1 [80]. Sequences were aligned using MAFFT v7.526 [81] and alignments were trimmed using trimAl in gappyout mode. Custom Python scripts were used for the rest of the analysis. A pairwise distance matrix of the 22381 sequences was computed using scipy v1.17.0 [82] with the Hamming distance as a metric. Agglomerative clustering was performed using scipy with the average linkage method and a threshold of 0.1 times the maximum distance was applied for the extraction of flat clusters. Minor clusters with less than a 100 sequences were excluded. Sequences comprised within each cluster were aligned using MAFFT. The alignments were trimmed using trimAl in gappyout mode and consensus sequences were computed using EMBOSS cons.

### De novo genome assembly

The ONT libraries for the wine subclade strains were assembled de novo using canu v2.2 [83], specifying a target size of 12 Mb. Contigs were assigned to chromosomes after alignment to the S288C reference genome using MUMmer v4.0.0 [84]. The assembly with the highest contiguity (YJM948) was selected for long read anchoring (below). The YJM948 assembly was masked using RepeatMasker v4.2.1 [85] in slow mode with a repeat library comprising Ty elements, the rDNA unit, subtelomeric repeats and TG_1-3_ repeats.

### Structural analysis of Y’ arrays

From the basecalled ONT libraries, the longest reads corresponding to 50X genome coverage (600 Mb, minimum length: 7992 bp, maximum length: 339654 bp, average length: 47444.1 bp) were subset using a custom Python script. To anchor the reads to chromosome ends, subset reads were aligned to the YJM948 canu assembly using minimap2 with preset lr:hq, and alignments of at least 20 kb in length were considered as candidate anchors. To annotate subtelomeric elements, subset reads were aligned to the masked S288C reference appended with the consensus sequence for each of the major Y’ types previously identified (see *Y’ types annotation and clustering*) using minimap2 with preset lr:hq. To annotate TG_1-3_ repeats, subset reads were aligned to the same reference as previously for subtelomeric elements using minimap2 with options -k15 -m10 -n1 -s30. The output of minimap2 is PAF files, which describe hits as sequence coordinates, similar to the most common annotation files like GFF and BED. Annotations in PAF format for chromosomal anchors, subtelomeric elements and TG_1-3_ repeats were combined and visualized using custom Python scripts. From these visualizations, the consensus structure of both haplotypes for each chromosome end was scored manually.

### Measurement of growth kinetics

Precultures were grown overnight in the same medium used for growth kinetics measurement and diluted 30-50X in fresh medium. 40 μL of cell dilutions were combined with 40 μL of medium with drug (if applicable) in a 384-well microplate (Corning cat. 3700). Optical density at 600 nm (OD600) was measured at 20 min intervals for 48 hrs at 30°C in a Cytation 3 microplate reader (Biotek Instruments Inc). For growth in permissive conditions, one replicate per strain was grown in YPD and SC. For cells carrying MoBY-ORF 2.0 *RAP1* or empty vector plasmids, six replicates per strain were grown in SC-MSG+G418. For dose-response curves in HU, one replicate per strain was grown in SC and SC supplemented with 11 concentrations (6-1024 mM) of HU (GoldBio cat. H-510-25). For the inhibition of haploid spores in 128 mM HU, 1-2 replicates per strain were grown in SC. For the inhibition of hemizygous *MAT* deletion strains in 128 mM HU, 3 replicates per strain were grown in SC. The analysis of growth curves was done using custom Python scripts. Maximum growth rates were extracted by fitting a modified logistic equation [86] to OD600 values. Inhibition was computed as the ratio of maximum rates in HU to SC. IC50 was derived by fitting the Hill equation to inhibition values, while IC25 and IC75 values were derived from the reciprocal of the fitted Hill equation.

### RNA extraction and transcriptome profiling

For strains YJM947, YJM955, YJM964 and YJM967, cells were grown to exponential phase at 30°C in 5 mL SC and SC supplemented with HU at the corresponding IC25, IC50 and IC75 concentrations. RNA was extracted using the YeaStar RNA Kit (Zymo Research cat. R1002). Transcriptome profiling libraries were prepared using the QuantSeq-Pool Sample-Barcoded 3’ mRNA-Seq Library Prep Kit (Lexogen cat. 139.96) and sequenced on a NovaSeq X Plus sequencer (Illumina) in paired-end 150 layout (Novogene). The library pool was demultiplexed using idemuxCPP v0.3.0 (Lexogen). Unique molecular identifiers (the first 10 bases of read 2) were extracted and added as read ID tags using cutadapt v5.1 [87] and UMI-tools v1.1.6 [88]. Reads were aligned to the masked S288C reference using Bowtie 2 v2.5.5 [89], and alignments were deduplicated using UMI-tools. A GTF annotation file was generated from the S288C GFF file (https://www.yeastgenome.org) using a custom Python script to include the 600 bp region at the 3’ of each gene, i.e. 250 bp upstream up to 350 bp downstream of the stop codon. Read counts per feature were extracted using Rsubread v2.24.0 [90] and differential expression analysis was run using DESeq2 [91] in custom R v4.5.2 scripts [92]. GO term enrichment analysis was performed using GOATOOLS v1.6.4 [93] in a custom Python script. Regulatory annotations were downloaded from AllianceMine and analyzed using a custom Python script. (https://www.alliancegenome.org/bluegenes/alliancemine, accessed on 4/6/2026)

### Genotype-phenotype associations

Biallelic variants within the wine subclade WT strains were selected in VCF format using bcftools and converted to raw genotype matrix format using PLINK. GWAS was run using rrBLUP v4.6.3 [94] with the maximum growth rate in YPD as a phenotype and VIII trisomy as a fixed effect. The threshold for statistical significance was defined as the 95^th^ percentile of the top score for each of 500 runs of the GWAS model with random permutation of the phenotype values. For the QTL analysis of spores, variants that are heterozygous in each parental background and homozygous in the spores were extracted using a custom Python script. Genetic distances (https://wiki.yeastgenome.org/index.php/Combined_Physical_and_Genetic_Maps_of_S._cerevisiae) were used to generate physical and genetic maps. QTL analysis against inhibition in 128 mM HU was run with r/qtl2 v0.40 [95] with the haploid cross type in a custom R script.

### MoBY-ORF 2.0 empty vector plasmid construction

Plasmids were extracted from the MoBY-ORF 2.0 yeast collection (Horizon cat. YSC11751). Cells grown in YPD+G418 were digested with zymolyase 20T at 37°C, lysed in alkaline buffer (SDS and NaOH) and neutralized with KOAc. DNA was precipitated in isopropanol and treated with RNAse A at 37°C. The MoBY-ORF 2.0 *RRM3* plasmid was used as a template to amplify the backbone with primers MHO_0030 and MHO_0031, and a 70 bp closing insert was generated with primers MHO_0001 and MHO_0002. A backbone:insert ratio of 1:5 was used in the assembly reaction. All plasmids were validated by long-read sequencing (Plasmidsaurus).

### Haploid spores collection

We generated a collection of haploid spores derived from each background of the wine subclade. We made deletion cassettes for *HO* with NatMX or HphMX cassettes with primers MHO_0021 and MHO_0022, using plasmids pAG25 and pAG32 as templates [96], respectively. WT competent cells were transformed with the cassettes and plated on YPD+Nat or YPD+Hyg. Hemizygous deletions were validated by PCR using primer pairs MHO_0023-MHO_0004/5 in 5’ and MHO_0024-MHO_0025/26 in 3’ for NatMX/HphMX. Cells of *HO* hemizygotes were sporulated on SPO plates for 3 days at room temperature. Tetrads were dissected on YPD plates using a MSM 300 semi-automatic dissection microscope (Singer Instruments) and incubated at room temperature for 3 days. Spore mating types were genotyped by PCR using primers MHO_0027, MHO_0028 and MHO_0029 [97].

### Hemizygous MAT deletion

Homozygous *HO* deletions were generated following the method described above, using the complementary deletion cassette and plating transformations on YPD+Hyg+Nat. KanMX deletion cassettes for the *MAT* locus were amplified from the pCAS plasmid [98] using primers MHO_0058 and MHO_0059. Transformations were plated on YPD+G418+Hyg+Nat. Hemizygous *MAT* deletions were validated by PCR using primer pairs MHO_0060-MHO_0003 and MHO_0061-MHO_0011. Mating types were genotyped by PCR as described above.

### RAD52-mNeonGreen tagging

The pCAS plasmid was mutagenized to include a gRNA sequence (GGCCAGGAAGCGTTTCAAGT) targeting the stop codon of *RAD52* using primers MHO_0077 and MHO_0078. A fragment containing the mNeonGreen fluorescent protein with 100 bp of flanking homology corresponding to the sequences upstream and downstream of the *RAD52* stop codon was synthesized (Twist Bioscience) and amplified with primers MHO_0075 and MHO_0076 to be used as donor DNA for CRISPR-Cas9 editing. Cotransformation of pCas-*RAD52* and the mNeonGreen fragment was done in *MAT**a*** spores of YJM947, YJM955, YJM964 and YJM967 that were selected for having the closest Y’ CN to their WT parent and plated on YPD+G418+Nat or YPD+G418+Hyg. Integration of the mNeonGreen tag was validated by PCR using primer pairs MHO_0079-MHO_0080 and MHO_0079-MHO_0081. Segregation of the pCAS plasmid after growing clones in YPD was monitored by spotting cells on YPD+G418 plates. Strains were mated with *MAT*α spores of the corresponding background that were selected for having the closest Y’ CN to their WT parent and transformed to switch the *HO* deletion cassette as described previously. Diploids were selected on YPD+Hyg+Nat.

### Cell cycle progression indicator plasmid construction

The cell cycle progression indicator plasmid (pRS31K_NLS-mTagBFP_*SPC42*-mCherry) was built as follows. To build plasmid pRS31K, the backbone of plasmid pRS31N [99] was amplified with primers MHO_0039 and MHO_0040 and the KanMX cassette was amplified from the pCAS plasmid using primers MHO_0041 and MHO_0042. A backbone:insert ratio of 1:2 was used in the assembly reaction. To build plasmid pRS31K_NLS-mTagBFP, pRS31K was digested with KpnI (NEB cat. R3142) and a fragment comprising the mTagBFP fluorescent protein with the N-terminal SV40 NLS peptide [100] was synthesized (Twist Bioscience) and amplified using primers MHO_0072 and MHO_0082. A backbone:insert ratio of 1:2 was used in the assembly reaction. To build plasmid pRS31K_NLS-mTagBFP_*SPC42*-mCherry, plasmid pRS31K_NLS-mTagBFP was digested with SacI (NEB cat. R3156) and a fragment comprising *SPC42* with the mNeonGreen fluorescent protein as a C-terminal tag was synthesized (Twist Bioscience) and amplified using primers MHO_0089 and MHO_0090. A backbone:insert ratio of 1:1 was used in the assembly reaction. All plasmids were validated by long-read sequencing (Plasmidsaurus).

### Cell imaging

Heterozygous *RAD52*-mNeonGreen strains were transformed with the cell cycle progression indicator plasmid and plated on YPD+G418+Hyg+Nat. Cells were grown in SC-MSG+G418 overnight, diluted 20 fold either in fresh SC-MSG+G418 medium or in SC-MSG+G418 supplemented with HU at the corresponding IC50 concentration, and grown for 3-4 hours at 30°C. Pads (∼0.3 mm thick) were prepared on glass slides with the corresponding medium supplemented with 1% agar. Cells in exponential phase were spotted on the pads, covered with no. 1.5 cover slips (Fisher Scientific cat. 12541013) and sealed with paraffin:mineral oil (1:2). Images were acquired on an Eclipse Ti-E microscope (Nikon) equipped with a 100X 1.45NA CFI Plan Apo Lambda objective (Nikon), a piezoelectric stage (Physik Instrumente), a CSU10 spinning disk confocal scanner unit (Yokogawa), 405, 488 and 561 nm lasers (Agilent Technologies), and an iXonUltra 897 EMCCD camera (Andor Technology). Z-stacks of 12 μm in 0.5 μm steps were acquired. Cells were checked for viability and cell cycle progression by monitoring the mTagBFP and mCherry channels. 25-40 fields were acquired for each genotype-condition combination, corresponding to 183-226 cells in each case. Maximum z-projection, level adjustment and channel merging were done using custom Python scripts.

## Declarations

### Ethics approval and consent to participate

Not applicable.

### Consent for publication

Not applicable.

### Availability of data and materials

Sequencing reads are available at NCBI under BioProject accession PRJNA1482565. Microscopy data are available upon request. Code and genome assemblies are available at https://github.com/mhenault1/subtelomeres_project. Exact command lines for all bioinformatics steps of the project are available in Supplemental File 1.

### Competing interests

The authors declare that they have no competing interests.

### Funding

L.R.H., M.H. and V.F. were supported by NIH K99/R00 grant (GM134193). M.H. was supported by postdoctoral fellowships from the National Sciences and Engineering Research Council of Canada (578152-2023) and from the Fonds de Recherche du Québec - Santé (331136). V.F. was supported by NIH T32 INCLUDE grant (GM136444-04S1). We acknowledge the NIH Cancer Center Support grant (P30CA046934) for support to the University of Colorado Cancer Center Flow Cytometry core.

### Authors’ contributions

L.R.H and M.H. designed the research. M.H. performed the experiments and data analysis. V.F. made the transcriptome profiling libraries. L.R.H. dissected tetrads for the project. M.H. and L.R.H. wrote the manuscript.

## Acknowledgements

We thank all members of the Heasley lab for discussions on the project, and Lisa M. Wood specifically for assistance with microscopy experiments. We thank Dr. John H. McCusker for giving us access to his collection of yeast isolates. We thank Amber Baldwin and Dr. Neelanjan Mukherjee for their resources and expertise with transcriptomics library preparation. We thank Dr. Jeffrey K. Moore for providing us training and access to his microscopy platform. We thank Dr. Michael A. McMurray for providing us access to his microplate reader. We thank Dr. Jay R. Hesselberth for providing us access to his ONT sequencer. We thank Dr. Lynn E. Heasley for providing us access to his computational resources. We thank Dr. Anne-Marie Dion-Côté for useful feedback on the manuscript.

---

**Figure S1.**
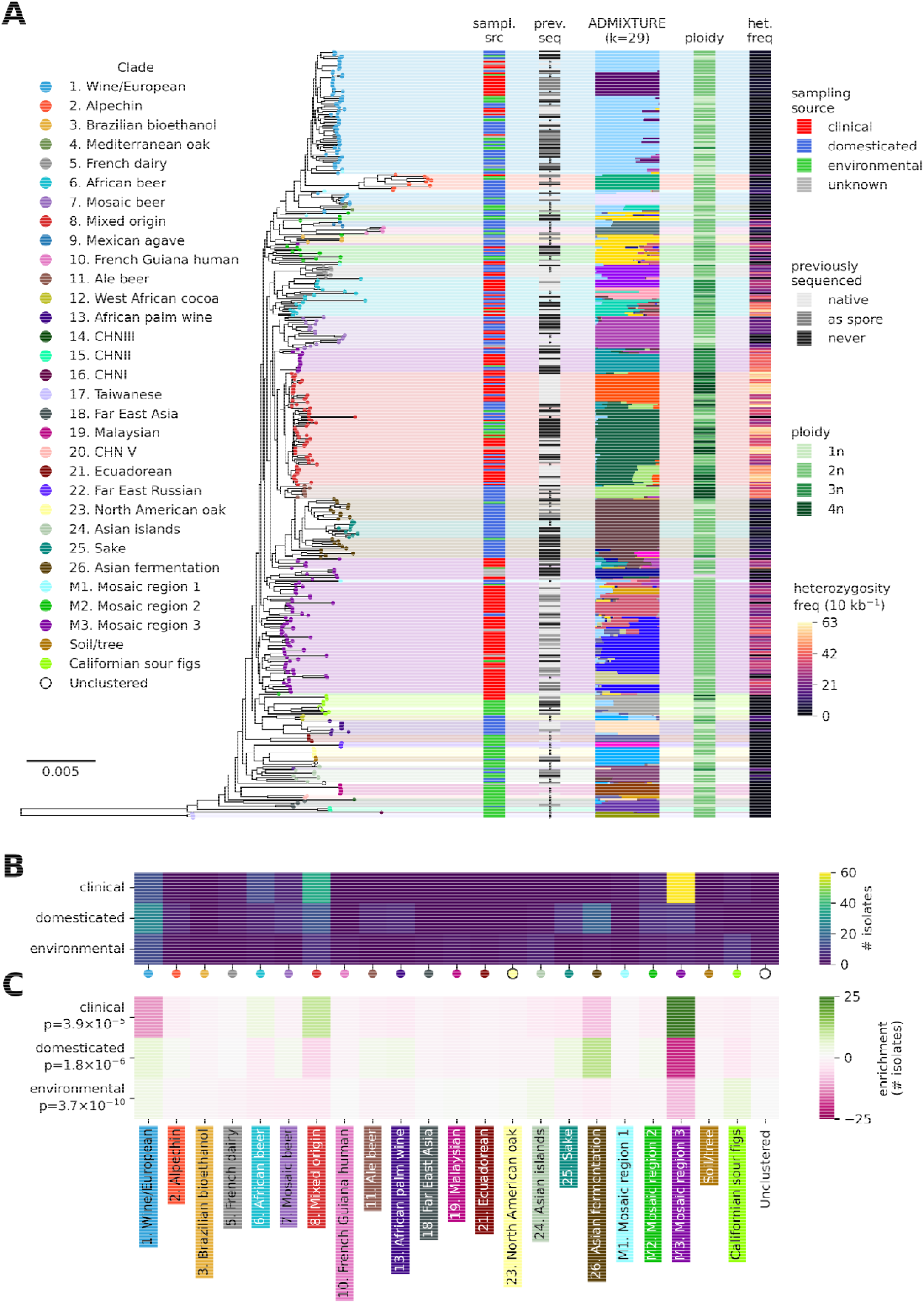
Population genomics analysis of the YJM collection yields a species-wide view of genetic variation. **A.** Maximum likelihood phylogenetic tree built from single nucleotide variants (SNVs) across 500 randomly sampled open reading frame (ORF) sequences from the nuclear genome. For each isolate, leaf color corresponds to the clade assignment. Annotations show (from left to right) sampling source; previous sequencing status (asterisks mark strains from the 1011GP included as reference); ancestry clusters defined by analysis with the ADMIXTURE software [78] at the optimum number of clusters (k=29); ploidy level measured by flow cytometry; and average genome-wide heterozygosity level, measured as heterozygous sites per 10 kb window. **B.** Counts of isolates per clade and sampling source. **C.** Deviation from the null expectation of equal representation across all phylogenetic groups. P-values are for chi-squared tests.

**Figure S2.**
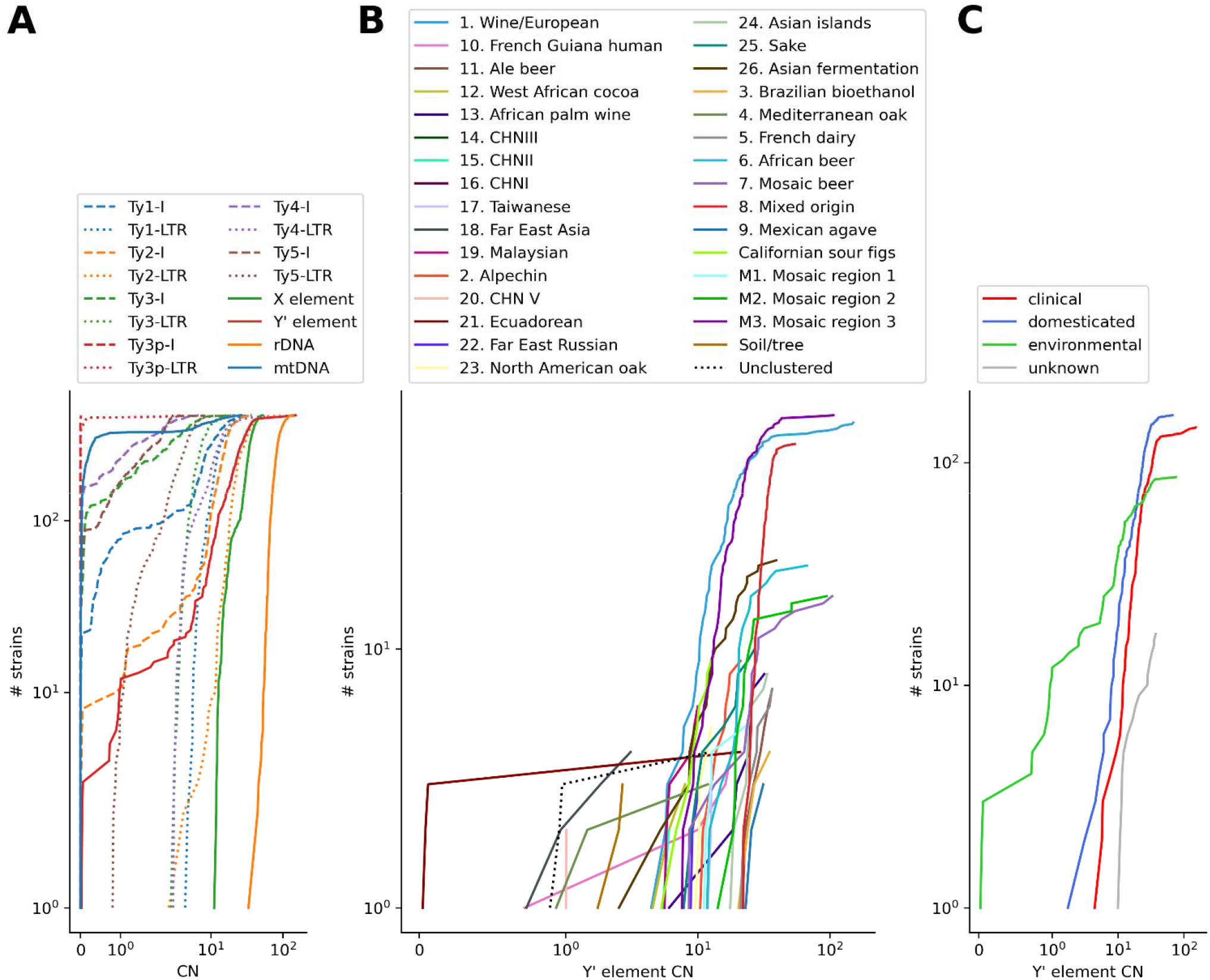
Species-wide CN distribution of genomic repeats. **A.** Distribution for all repetitive element families. **B.** Distributions of Y’ element CN per clade. **C.** Distributions of Y’ element CN per sampling source. Distributions are shown as empirical cumulative distributions, with both axes in symlog scale with a linear threshold of 1.

**Figure S3.**
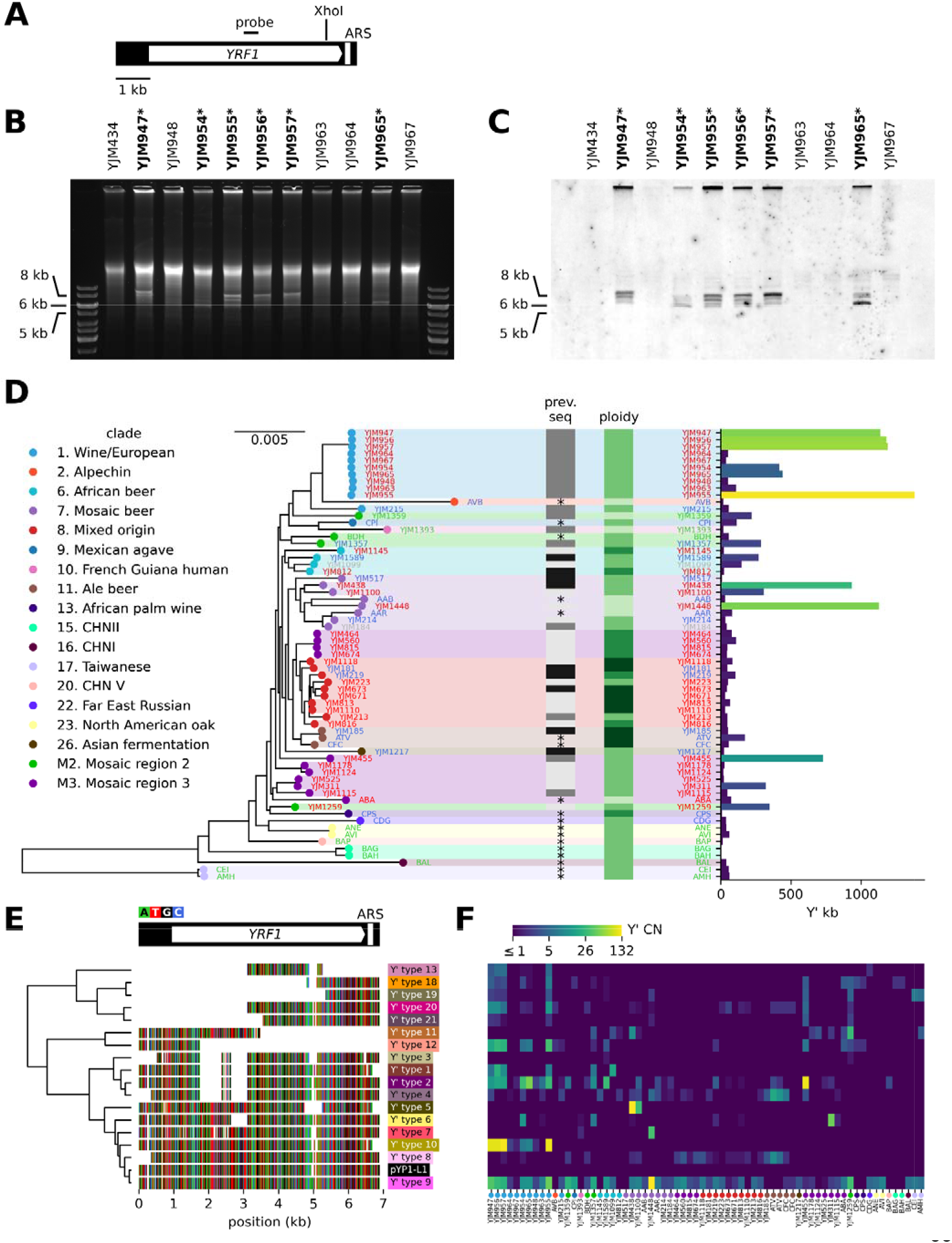
Species-wide Y’ expansions are driven by multiple Y’ sequence types. **A.** Annotation of the canonical Y’ element, including the location of the probe and the single XhoI restriction site commonly used for Southern analysis of Y’ elements. **B.** Ethidium bromide (EtBr) staining of genomic DNA digested with XhoI. Strains with Y’ amplifications are labelled with asterisks. YJM434 is included as an outgroup. **C.** Hybridization with Y’ probe labelled with AlexaFluor 488. **D.** Subset of the species phylogenetic tree in Fig. S1 showing the YJM strains selected for Oxford Nanopore long-read sequencing and the reference strains for which haploid long-read assemblies are available. Y’ CN estimates shown are derived from raw read and assembly annotations, respectively. Strain names are colored by sampling source using the color code in Fig. S1. Previous sequencing and ploidy color codes are the same as in Fig. S1. **E.** Multiple sequence alignment of the 17 consensus sequences of Y’ types. Annotations at the top show the structure of the canonical Y’ element from [25] (pYP1-L1), including the autonomously replicating sequence (ARS) and the approximate coordinates of the main helicase and serine/threonine-rich domains within *YRF1*. **F.** CN distribution across Y’ types for the strains shown in panel A. CN color mapping is on a logarithmic scale. Duplicate labels on the X axis correspond to collapsed and phased assemblies for polyploid strains [33].

**Figure S4.**
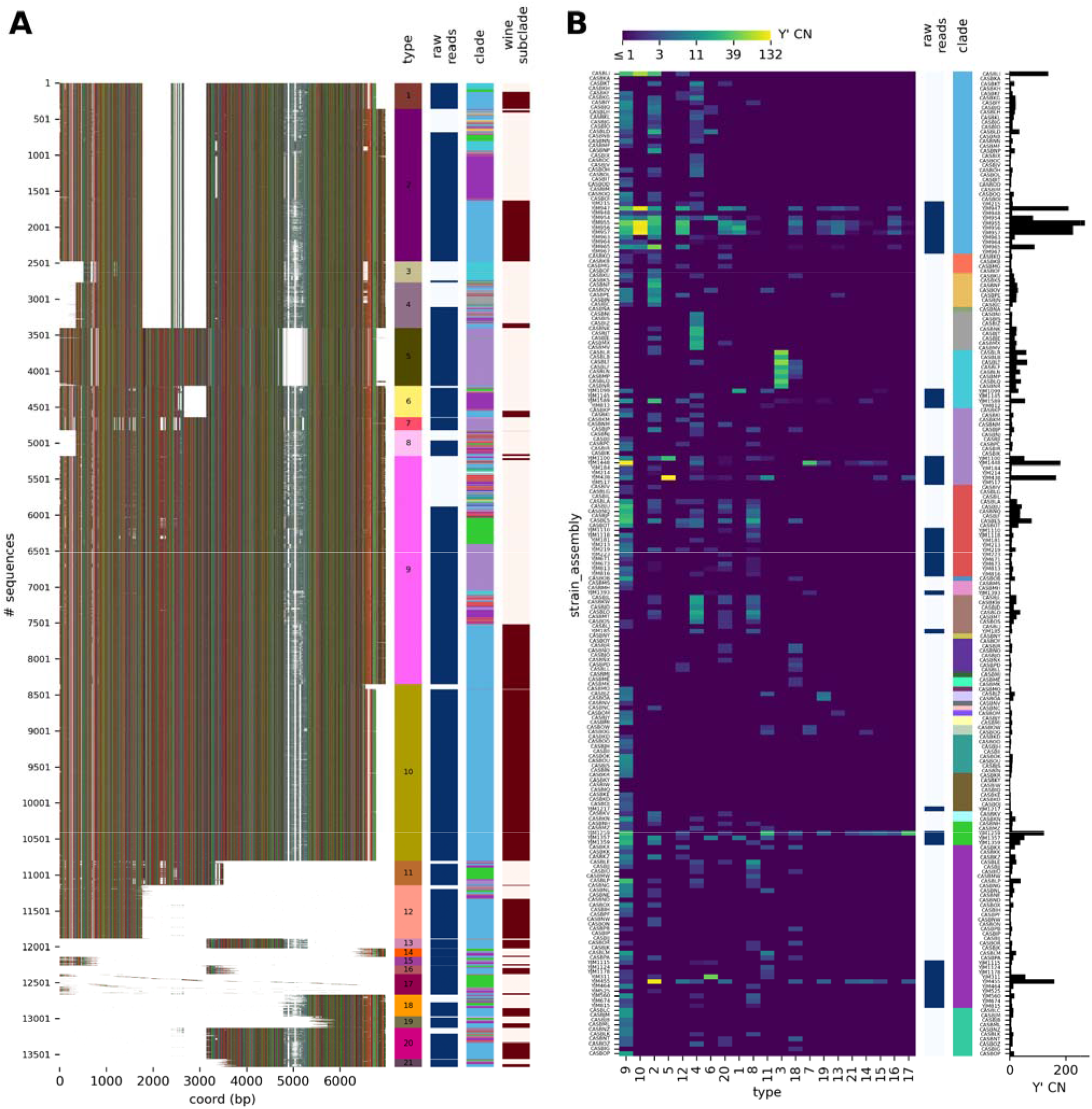
Clustering of the species-wide repertoire of Y’ sequences. **A.** Multiple sequence alignment (MSA) and clustering of 13678 Y’ sequences. Clusters comprising less than 100 sequences were excluded. Annotation columns at the right of the MSA show sequence type identified by agglomerative clustering; sequence source (dark: annotation of raw reads, light: annotation of de novo assemblies [33]); clade of the source strain; and whether the source strain is in the wine subclade (dark: true, light: false). **B.** Y’ CN profile by type across the raw reads (dark) and genome assemblies (light). Total estimated CN is shown at the right.

**Figure S5.**
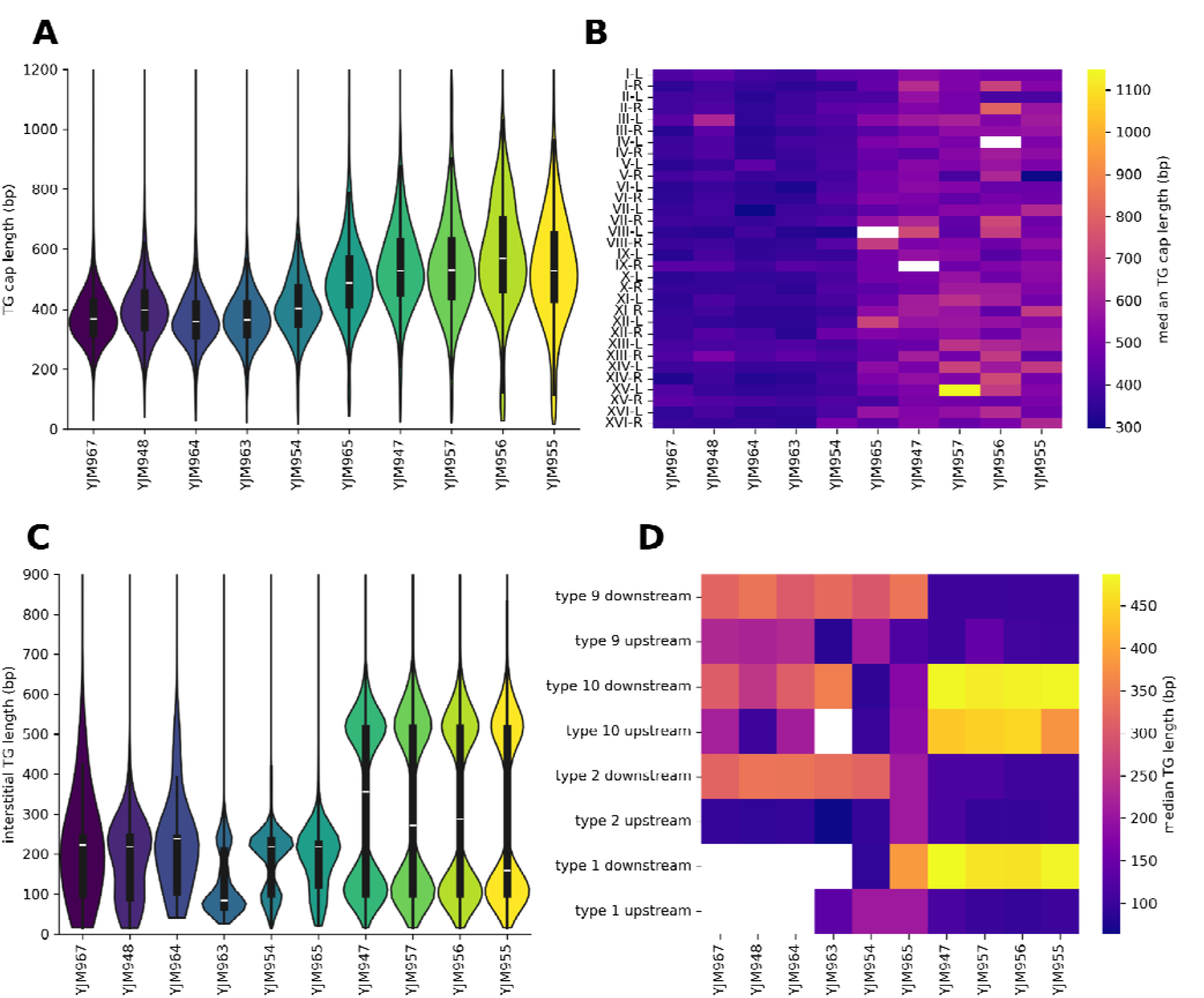
TG_1-3_ repeat length variation is associated with Y’ contents. **A.** Distributions of TG_1-3_ repeats at telomere caps. Strains are ordered by increasing Y’ CN. **B.** Median telomere cap length at individual chromosome ends. **C.** Distributions of interstitial TG_1-3_ repeats within Y’ tandem arrays. **D.** Median TG_1-3_ repeat length for TG_1-3_ repeats found immediately adjacent to the most common Y’ types. For each Y’ type, the median length is shown for TG_1-3_ repeats located either directly upstream (towards the centromere) or directly downstream (towards the telomere cap, including the telomere cap) of the Y’ element.

**Figure S6.**
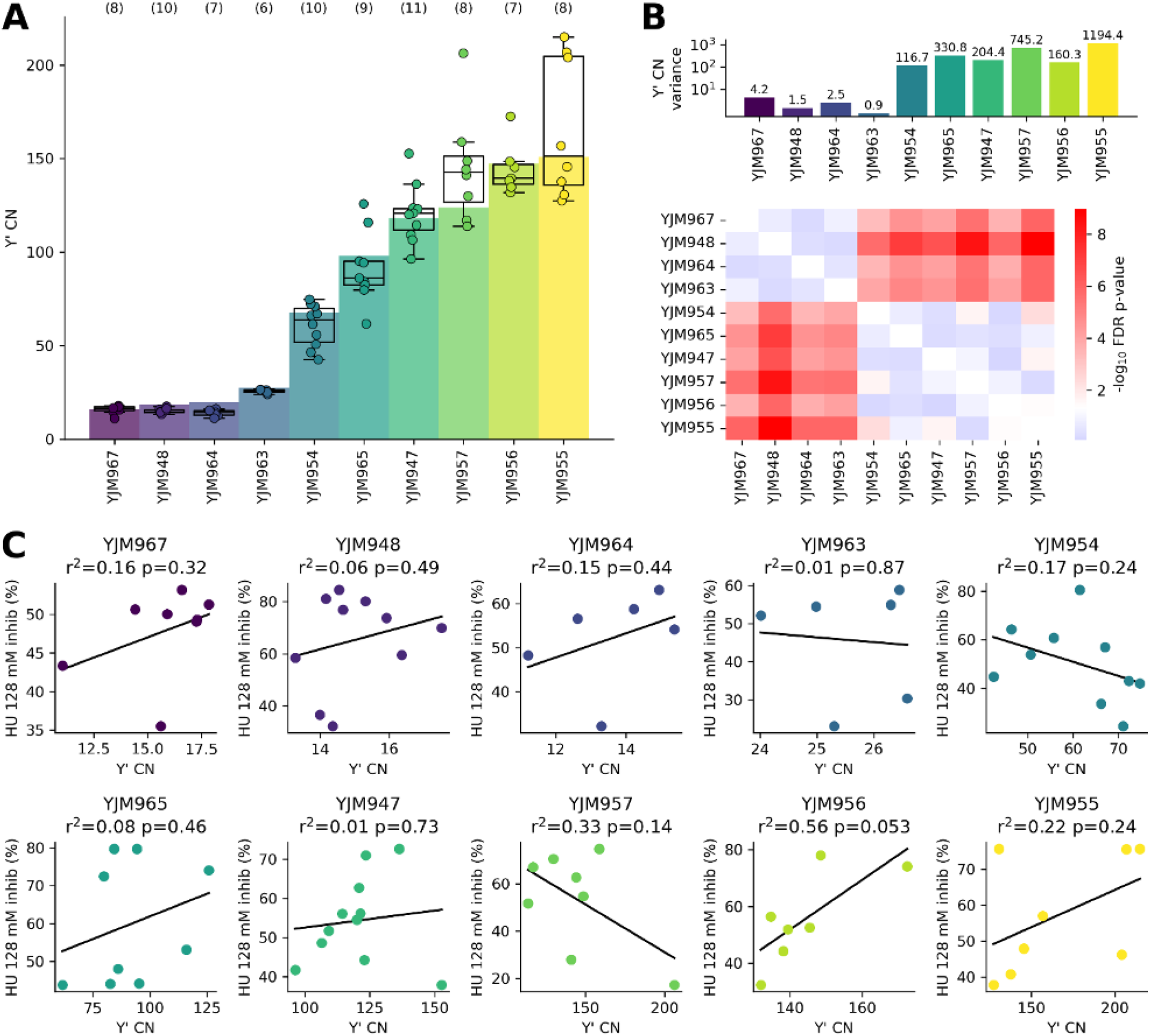
Y’ CN variation in haploid spores of the wine subclade. **A.** Distribution of Y’ CN for haploid spores of each background. Bars indicate the CN of the corresponding WT parent. **B.** Variance of CN for the spores of each background (top) and pairwise Bartlett’s tests for the difference of variance between CN distributions of spores from each background. The color map shows the FDR-adjusted *p*-values, where red hues correspond to values below *p*-value=0.05. **C.** Correlation between Y’ CN of the haploid spores and sensitivity to HU, measured as the ratio between growth at 128 and 0 mM HU.

**Figure S7.**
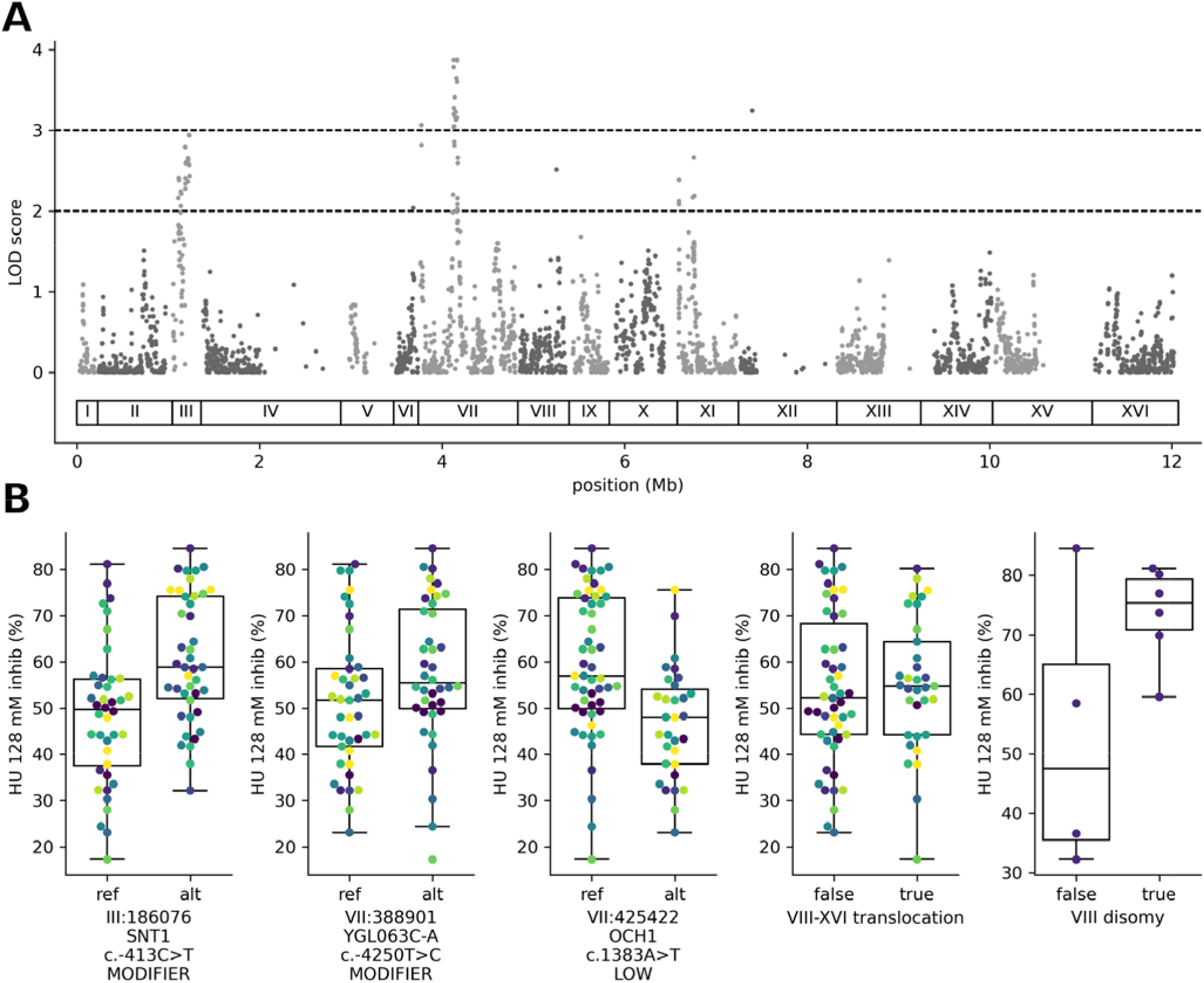
Correlation of spore genotypes with HU sensitivity. **A.** Quantitative trait loci (QTL) analysis for HU sensitivity. Two significance levels corresponding to LOD scores of 2 and 3 are shown as dashed lines. **B.** Distributions of growth inhibition in 128 mM HU as a function of the segregation of 3 significant QTL SNVs, the VIII-XVI translocation that is heterozygous in the whole wine subclade [101], and disomy of VIII in YJM948 spores. In the latter case, the increase in HU sensitivity in disomic spores is marginally significant (Student’s *t*-test, *p*-value: 0.08).

**Figure S8.**
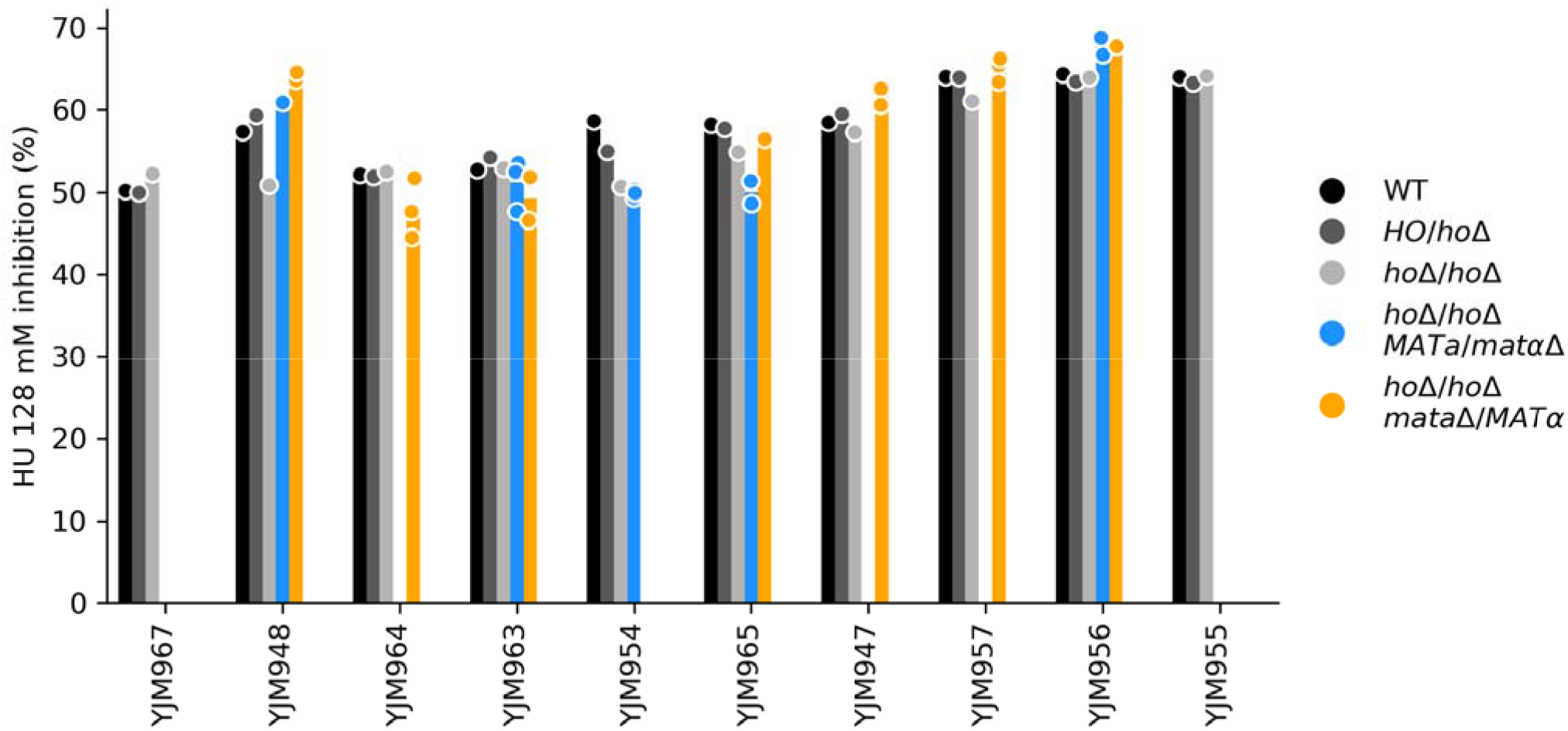
HU sensitivity in diploid backgrounds of the wine subclade as a function of *HO* and *MAT* genotype. Percent inhibition in 128 mM HU is shown for WT strains and sequentially derived hemizygous *HO* deletion strains, homozygous *HO* deletion strains and hemizygous *MAT* deletions, where applicable. The *HO* locus encodes an endonuclease responsible for triggering mating-type switching in cells of *MAT**a** or MAT*α identity [102].

**Figure S9.**
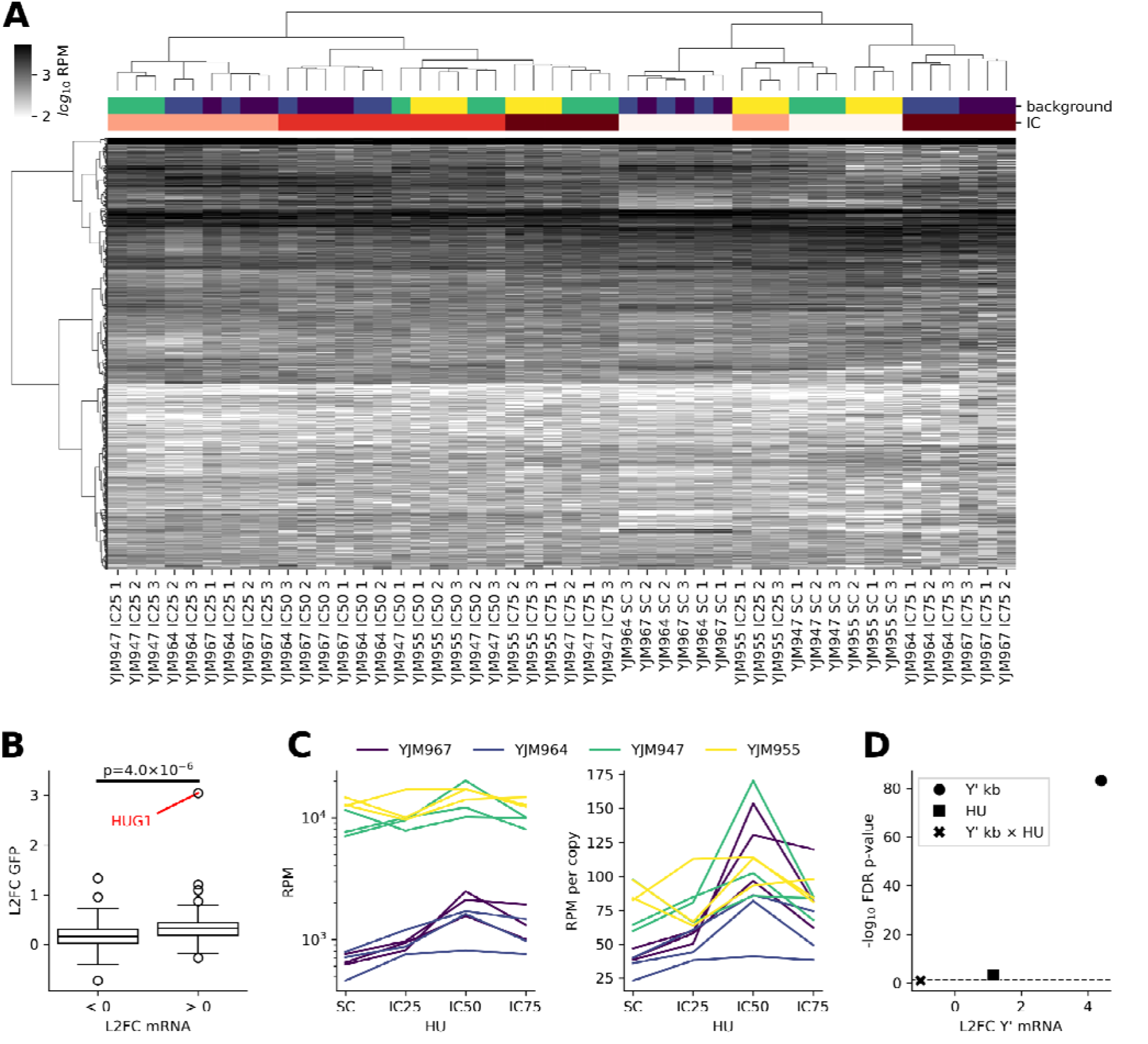
Transcriptome profiling of the wine subclade as a function of Y’ CN and HU exposure. **A.** Transcript levels shown as log_10_ RPM for the three replicate libraries of each condition. Genes (rows) with no zero counts are included. Color codes (columns) show the background (see Fig. 1F) and HU inhibition level (white: SC, light red: IC25, medium red: IC50, dark red: IC75). **B.** Expression level of genes measured by the yeast GFP collection in 200 mM HU [50] as a function of DE level for the HU contrast. P-value for Student’s *t*-test is shown. **C.** Transcript levels for the Y’ element as a function of HU exposure level, either raw (left) or normalized by Y’ CN (right). **D.** Fit of the DE model for the Y’ element transcript level as a function of Y’ abundance, HU exposure and their interaction. The dashed line indicates a significance threshold of 0.05.

**Figure S10.**
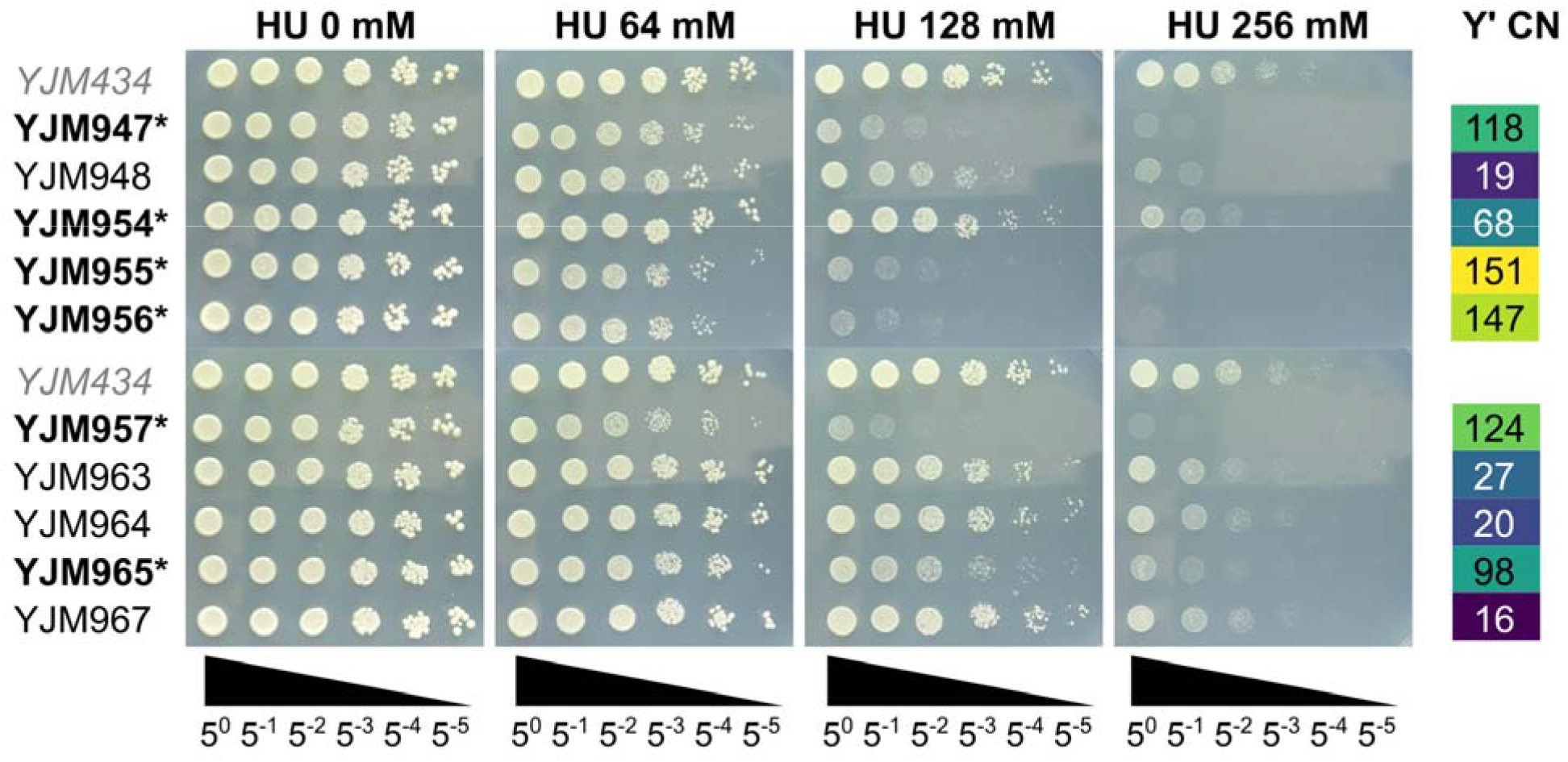
Spot assays of the wine subclade on HU media. Precultures of equal OD600 were diluted five-fold and spotted on plates containing increasing concentrations of HU. Strains with Y’ amplifications are shown in bold, with the corresponding Y’ CN at the right. YJM434 is included as an outgroup.

## Notes

https://github.com/mhenault1/subtelomeres_project

### Competing Interest Statement

The authors have declared no competing interest.

### Summary of Updates

Main body text has received minor edits to clarify specific experimental approaches. Figures 4, S2, and S6 have been updated.

